# A marine sponge-associated mycobacterium closely related to *Mycobacterium tuberculosis*

**DOI:** 10.1101/2024.01.23.576949

**Authors:** Sacha J. Pidot, Stephan Klatt, Louis S. Ates, Wafa Frigui, Fadel Sayes, Laleh Majlessi, Hiroshi Izumi, Ian R. Monk, Jessica L. Porter, Vicki Bennett-Wood, Torsten Seemann, Ashley Otter, George Taiaroa, Gregory M. Cook, Nicholas West, Nicholas J. Tobias, John A. Fuerst, Michael D. Stutz, Marc Pellegrini, Malcolm McConville, Roland Brosch, Timothy P. Stinear

## Abstract

Reconstructing the evolutionary origins of *Mycobacterium tuberculosis*, the causative agent of human tuberculosis, has helped identify bacterial factors that have led to the tubercle bacillus becoming such a formidable human pathogen. Here we report the discovery and detailed characterization of an exceedingly slow growing mycobacterium that is closely related to *M. tuberculosis* for which we have proposed the species name *Mycobacterium spongiae* sp. nov., (strain ID: FSD4b-SM). The bacterium was isolated from a marine sponge, taken from the waters of the Great Barrier Reef in Queensland, Australia. Comparative genomics revealed that, after the opportunistic human pathogen *Mycobacterium decipiens*, *M. spongiae* is the most closely related species to the *M. tuberculosis* complex reported to date, with 80% shared average nucleotide identity and extensive conservation of key *M. tuberculosis* virulence factors, including intact ESX secretion systems and associated effectors. Proteomic and lipidomic analyses showed that these conserved systems are functional in FSD4b-SM, but that it also produces cell wall lipids not previously reported in mycobacteria. We investigated the virulence potential of FSD4b-SM in mice and found that, while the bacteria persist in lungs for 56 days after intranasal infection, no overt pathology was detected. The similarities with *M. tuberculosis*, together with its lack of virulence, motivated us to investigate the potential of FSD4b-SM as a vaccine strain and as a genetic donor of the ESX-1 genetic locus to improve BCG immunogenicity. However, neither of these approaches resulted in superior protection against *M. tuberculosis* challenge compared to BCG vaccination alone. The discovery of *M. spongiae* adds to our understanding of the emergence of the *M. tuberculosis* complex and it will be another useful resource to refine our understanding of the factors that shaped the evolution and pathogenesis of *M. tuberculosis*.

## Introduction

*M. tuberculosis,* the causative agent of human tuberculosis (TB), is the leading bacterial cause of mortality and morbidity worldwide and is responsible for approximately 1.5 million deaths per year (1). Tuberculosis has affected humans since at least the neolithic expansion of humans across the continents. Despite the wealth of molecular evidence explaining the evolution of mycobacteria that cause tuberculosis in humans and other mammals (the *Mycobacterium canettii* clade and *M. tuberculosis* complex, MTBC), the origins of this complex and their differentiation from other mycobacteria are only beginning to be understood.

Several environmental mycobacteria have also been noted as close ancestors of the MTBC, including *Mycobacterium marinum*, a fish and human pathogen (2), and *Mycobacterium kansasii* (3), although neither of these mycobacteria have been seen to transmit between humans and they have significantly larger genomes than *M. tuberculosis.* Recent genomic analyses have identified some other opportunistic human pathogens, such as *Mycobacterium riyhadense*, *Mycobacterium lacus, Mycobacterium shinjukense* and *Mycobacterium decipiens*, that share many features of host-adaptation with *M. tuberculosis* (4). However, these closely related, slow growing non-tuberculous mycobacteria (NTM) bacteria differ in many aspects from tuberculosis-causing mycobacteria. These studies suggest that there are likely other taxa to discover that can aid our understanding of *M. tuberculosis* evolution from a generalist mycobacterium into a highly virulent, specialist human pathogen.

Marine sponges are known to house a large and diverse repertoire of bacteria, as attested by recent efforts to catalogue the microbiome of these animals from around the world (5–7). These studies have shown that Actinobacteria are one of the largest phyla within these microbial communities. As part of efforts to identify and catalogue Actinobacterial symbionts from marine sponges on the Great Barrier Reef in Australia, and to identify possible target species of anti-mycobacterial rifamycins produced by *Salinospora* sponge symbionts, a mycobacterial isolate named FSD4b-SM was isolated from a *Fascaplysinopsis reticulata* sponge at a depth of 25m (8). Initial investigations of this strain showed that it was closely related to the MTBC by conserved gene amplicon sequencing (8).

Here, we sought to better understand the genetic and functional relationships between FSD4b-SM and the MTBC through genomic, proteomic and lipidomic analyses. Our research establishes FSD4b-SM as the most closely related marine organism to the MTBC, assesses its virulence potential, and its prospects for use in TB vaccine development.

## Materials and Methods

### Culture conditions

*M. spongiae* was grown in simplified marine broth (5 g/L peptone, 1 g/L yeast extract, 33 g/L artificial sea salt). For growth on plates, simplified marine broth was supplemented with 10 g/L bacteriological agar (Difco). Cultures were incubated at 28°C for 2 to 3 months without shaking. A list of strains and plasmids used in this study can be found in Table S1.

### Electron microscopy

Transmission electron microscopy was performed by first washing *M. spongiae* FSD4b-SM cells from 3 month old cultures in PBS and pelleting by centrifugation at 10,000 x *g* for 5 min. Cells were then resuspended in fixation buffer (2.5% glutaraldehyde in 0.1 M sodium cacodylate) and incubated for 2 h at RT. Cells were then pelleted by centrifugation and washed twice with 0.1 M sodium cacodylate before post-fixation in 1% osmium tetroxide for 2 h at RT. Cell pellets were then washed in dH_2_O, left overnight at 4°C in 0.3% uranyl acetate and then rinsed with dH_2_O before being dehydrated using a graded series of acetone. Samples were then infiltrated and embedded with EPON resin. Sections (70–80 nm thick) were cut and stained with uranyl acetate and lead citrate before being viewed under a Phillips CM120 transmission electron microscope at 120 Kv.

### Genome sequencing

High molecular weight genomic DNA was prepared using the DNeasy Blood and Tissue kit (QIAgen), according to the manufacturer’s instruction for Gram positive bacteria. A complete FSD4b-SM genome sequence was generated using a combination of PacBio and Illumina sequencing. For sequencing on the PacBio RSII, extracted DNA was prepared using the Template Prep Kit 1.0 (PacBio) and following adapter ligation DNA was size selected using a BluePippin system (Sage Biosciences) with a 8 kb cut-off. Adapter-ligated, circularised DNA was loaded onto a single SMRT cell at 0.2 nM and sequence data were captured with a 6 h movie time. PacBio sequencing data was assembled using HGAP3, as implemented in the SMRT Portal (PacBio). The resulting genome was polished three times using Quiver (PacBio) before being error corrected with Illumina reads using Snippy v3.2 (https://github.com/tseemann/snippy). For Illumina sequencing, DNA libraries were created using the Nextera XT DNA preparation kit (Illumina) and whole genome sequencing was performed on the NextSeq platform (Illumina) with 2 x 150bp paired-end chemistry. A sequencing depth of >50× was targeted and the reads were used for error correcting the PacBio-assembled genome, as outlined above. The final 5,581,157 bp genome sequence was annotated using the NCBI Prokaryotic Genome Annotation Pipeline (PGAP) and assigned Genbank accession number CP046600.

### Bioinformatics

Pairwise whole genome average nucleotide identity (ANI) was calculated using *fastANI* (https://github.com/ParBLiSS/FastANI) and *ANIclustermap* (https://github.com/moshi4/ANIclustermap) (9). Core genome and ortholog comparisons were performed using *bcgTree* (10) and *Roary* (11). Phylogenies were inferred using *iqtree* (12) using the the protein sequence alignment file output of *bcgTree*, with 1000 bootstrap replicates and the JTT model of amino acid substitution. A list of the mycobacterial genomes used for comparisons can be found in Table S2. Individual protein homology searches were performed using BLAST, as implemented at NCBI (https://www.ncbi.nlm.nih.gov/), with multiple amino acid sequence alignments performed with ClustalW (13) and phylogenetic trees built using the Geneious tree builder (Geneious v9.1) (https://www.geneious.com). Analysis and alignment of ESX loci with *cblaster* (14) and *clinker* (15) was performed using the online *CompArative GEne Cluster Analysis* Toolbox (cagecat.bioinformatics.nl). Alignments of other loci were performed with *clinker* (15) at cagecat.bioinformatics.nl. PE/PPE proteins were identified through homology searching using BLAST and via annotation describing the protein as either a PE or PPE family member. All putative *M. spongiae* PE/PPE proteins were then aligned against known PE/PPE proteins and investigated for the presence of key domains/signature sequences according to criteria in (16). Analysis of specialised metabolism was performed with antiSMASH (17).

### Extraction of and LC-MS analysis of mycobacterial lipids

Mycobacterial lipids were extracted and analysed as previously described (18). In brief, cell pellets were extracted in 20 volumes of chloroform/methanol (2:1, v/v) followed by chloroform/methanol/water (1:2:0.8, v/v/v). Insoluble material was removed by centrifugation, extracts were dried under nitrogen and subjected to biphasic partitioning in 1-butanol and water (2:1, v/v). The organic phase was dried and lipids were resuspended in water-saturated 1-butanol. Lipid extracts were separated on an Agilent 1290 Infinity LC System (Agilent Technologies) using a Kinetex C 18 column (Phenomenex; 2.6 µm EVO C18 100Å) and eluted by using the following binary solvent system: mobile phase A (ACN:H2O (60:40, v/v) with 10 mM ammonium formate) and mobile phase B (IPA:ACN (90:10, vol/vol), with 10 mM ammonium formate), with a 30 min gradient program. Eluted lipids were detected using a 6550 iFunnel QTOF LC/MS system (Agilent Technologies) with the same parameters as previously described (18). Lipids were identified in positive ionization mode by accurate mass, fragmentation pattern, retention time and retention time order (different lipid groups and different saturation levels show an elution time pattern and a relation to each other). MS-DIAL (Version 2.06; MS/MS data) was used for manual lipid annotation.

### Proteomics

Samples for proteomics were prepared from FSD4B-SM cultures using the SP3 method (19). Briefly, cells were lysed using buffer containing 50 mM HEPES, pH 8, 1% (wt/vol) SDS, 1% (vol/vol) Triton X-100, 1% (vol/vol) NP-40, 1% (vol/vol) Tween 20, 1% (wt/vol) deoxycholate, 5 mM EDTA, 50 mM NaCl, 1% (vol/vol) glycerol and beat beating for 6 x 30s in a Precellys 24 at speed 6.5. Insoluble material was removed by centrifugation at 20,000 x g for 10 min and the supernatant was transferred to a fresh tube. Protein concentration was measured using the BCA assay kit (Thermo Scientific) with bovine serum albumin as a standard. 10 µg protein was bound to SeraMag SpeedBeads Carobxylate-modified [E3] (Cytiva) and were digested overnight at 37°C with a 1:50 trypsin:protein ratio. Tryptic peptides were recovered and were cleaned up through SDB-RPS resin prior to submission for mass spectrometry.

The purified peptide samples were analysed via nano liquid chromatography coupled to tandem mass spectrometry (LC-MS/MS) at the University of Melbourne Mass Spectrometry and Proteomics Facility, using an Orbitrap Eclipse Tribrid Mass Spectrometer (Thermo Fisher Scientific, USA) equipped with a nano ESI interface coupled to an Ultimate 3000 nano HPLC (Thermo Fisher Scientific, USA). Peptides were separated using an Acclaim PepMap RSLC analytical column (C18, 100 Å, 75 μm × 50 cm, Thermo Fisher Scientific, USA) and Acclaim PepMap trap column (75 μm × 2 cm, C18, 100 Å). The enrichment column was injected with the tryptic peptides (3 µL) at an isocratic flow of 5 μL/min of 2% v/v CH3CN containing 0.05% v/v aqueous trifluoroacetic acid for 6 min, applied before the enrichment column was switched in-line with the analytical column. The eluents were 0.1 % v/v aqueous formic acid and 5 % v/v dimethyl sulfoxide (DMSO) (solvent A) and 0.1 % v/v formic acid and 5 % DMSO in acetonitrile (solvent B). The gradient was at 300 nL min-1 from (i) 0–6 min, 3 % B; (ii) 6–35 min, 3–23 % B; (iii) 35-45 min, 23-40 % B; (iv) 45-50 min, 40-80 % B; (v) 50-55 min, 80–80 % B; (vi) 55-55.1 min, 80-3 % B; (vii) 55.1-65 min, 3-3 % B. The column oven was maintained at 50 °C throughout the analysis. The Eclipse Orbitrap mass spectrometer was operated in the data-dependent mode, wherein full MS1 spectra were acquired in a positive mode over the range of m/z 375-1500, with spray voltage at 1.9kV, source temperature at 275 °C, MS1 at 120,000 resolution and normalized AGC target of 100 % and maximum ion injection time of 50 ms. The top 3 second method was used and selecting peptide ions with charge states of ≥ 2-7 and intensity thresholds of ≥ 5E4 were isolated for MS/MS. The isolation window was set at 1.6 m/z, and precursors were fragmented using higher energy C-trap dissociation (HCD) at a normalised collision energy of 30 %, a resolution of 15,000, a normalized AGC target of 100% and automated IT time.

### Protein fractionation and Western blotting of M. spongiae

*M. spongiae* FSD4B-SM liquid preculture of 25 ml was grown until an OD600 of 0.15 was reached. At this point the complete culture was added to fresh simplified marine broth to a total volume of 100 ml. The cultures were incubated for 14 days in standing conditions at 30°C. After the 14 days of incubation, the culture had reached a density of 0.203 OD_600_/ml. The culture was centrifuged in 2 x 50 ml falcon tubes at 5000 rpm for 10 minutes. The supernatant was taken and centrifuged again at 5000 rpm, after which the remaining supernatant (2 x 45ml) was collected and concentrated over a 3 kDa Amicon filter (Millipore). The residue was suspended in a volume of 900 µl. Pelleted cells after the first centrifugation step were washed once with PBS and resuspended in solubilisation buffer and boiled at 95°C for 15 minutes. An equivalent of 0.3 OD_600_ units of culture was loaded in each lane. *Mycobacterium tuberculosis* CDC1551 samples were loaded as a comparator. CDC1551 samples were taken from previously published protein fractions (20). SDS-Page total protein content was imaged using standard Coomassie brilliant blue staining with Kaleidoscope molecular weight ladder (Biorad). Antibodies tested to visualize *M. spongiae* proteins included Anti-PGRS 7C4.1F7 (21) (Clone 7C4.1F7 was a kind gift from Michael J. Brennan, USA); polyclonal anti-SigA (Kind gift from I. Rosenkrands, Denmark); monoclonal ESAT-6 (hyb76-8) (Harboe et al 1998); anti-GroEL2 antibody CS-44 (Kind gift from J. Belisle, Colorado State University, Fort Collins, CO); Rabbit polyclonal anti-EsxN (rMTb9.9A)(22).

### TAR cloning of the M. spongiae ESX-1 region

To design primers for amplification of the FSD4b-SM ESX-1 locus, the region was divided into 8 theoretical fragments of approximately equivalent length. Primers were then designed so that each PCR fragment would overlap by at least 30bp of DNA (see Table S3 for primer sequences). DNA fragments were amplified using Phusion DNA polymerase (NEB) in reactions containing 10% DMSO and using GC buffer. The program for amplification was: 1 cycle of 95°C for 5’ then 30 cycles of 95°C for 10s, 72°C for 2 min 30s followed by a final extension at 72°C for 7 minutes. PCR products were cleaned up using AMPure XP beads (Beckman Coulter). The GeneArt® High-Order Genetic Assembly kit (Thermo Fisher Scientific) was used to assemble the PCR products into the vector pYES1L by transformation into *Saccharomyces cerevisiae* MaV203 cells. The manufacturers protocols were followed for all steps. Recovered pYES1L:ESX-1^FSD4b-SM^ DNA was sequenced using Illumina sequencing to confirm the correct sequence. The 33.3 kb ESX-1^FSD4b-SM^ region from pYES1L:ESX-1^FSD4b-SM^ was then subcloned by digestion with *Sbf*I and gel extraction followed by ligation into the mycobacteria-*E. coli* shuttle vector pYUB412 that had been digested with *Sbf*I. The resulting ligation mixture was transformed by electroporation into *E. coli* DH10B and colonies were screened by PCR for the presence of the ESX-1^FSD4b-SM^ region. Cloning of the full-length ESX-1^FSD4b-SM^ region was confirmed by restriction digestion. The resulting pYUB412:ESX-1^FSD4b-SM^ plasmid was transformed into *M. bovis* BCG by electroporation according to standard protocols (23).

### qPCR to estimate M. spongiae concentration in mouse tissues

A standard TaqMan assay was designed for quantitative, specific detection of *M. spongiae* in mouse tissue specimens. A PCR amplicon was designed spanning a 64bp region of F6B93_05840, a CDS encoding a hypothetical aquaporin protein. The primer sequences were: FSD4b_05840-F 5’-ACGTCAGGCTTGATGCTCTC-3’ and FSD4b_05840-R 5’- GCGCTACCAGATAGACCCAG - 3’. The internal probe sequence was FSD4b_05840-P: 5’- [6FAM]CGGGTTTTTCTCGTGGAAGT[BHQ1] - 3’. The qPCR was performed as described (24) using 2x SensiFast mastermix (Bioline) with primers and probes to a final concentration in 25uL reaction volume of 0.32uM (primers) and 0.16uM (probe) respectively. Each reaction included internal positive control reagents and DNA template with 2uL volume of sample template DNA. PCR cycles included 1x 95oC 5min followed by 45x 95oC for 10s and then 60°C for 20s. PCR was performed in a LC480 Lightcycler (Roche). A standard curve was prepared using 10-fold serial, replicate dilutions of *M. spongiae* purified genomic DNA. Genomic DNA concentrations were measured using fluorimetry (Qubit, Thermofisher).

### Mouse vaccination and infectious challenge

Wild-type C57BL/6 mice were bred and maintained at The Walter and Eliza Hall Institute of Medical Research Animal Facility. Intranasal and subcutaneous vaccinations were performed in 50 µL volumes in PBS containing either 10^4^ CFU of live *M. spongiae*, or 5 x 10^4^ CFU of the WT or BCG::ESX-1^FSD4b-SM^ modified *M. bovis* BCG strains (the same dose was given for both intranasal and subcutaneous vaccination.

For *M. tuberculosis* infectious challenge, mice were infected with 50-200 colony-forming units (CFU) of *M. tuberculosis* H37Rv by aerosol using a whole-body Inhalation Exposure System (Glas-Col) four month after vaccination, as described (25). A bacterial suspension containing ∼2.5×10^8^ CFU in 6mL was aerosolized over a period of 45 min. Mice were euthanized four weeks post-infection by CO_2_ asphyxiation. Spleens and the right lung lobes were aseptically harvested and homogenized with steel beads in PBS+0.05% Tween-80 using a Bullet Blender (Next Advance) at setting #6 for 3 min (spleens) or #8 for 5 min (lungs). Tissue homogenates for counting *M. tuberculosis* were serially diluted and spread on 7H11 agar plates (BD Biosciences) supplemented with 0.5% glycerol and 10% (v/v) oleic-albumin-dextrose-catalase supplements (50 g/l BSA, 20 g/l dextrose, 0.04 g/l catalase and 0.5 g/l oleic acid [Sigma-Aldrich]). Plates were incubated at 37°C for 3 weeks before counting. *M. spongiae* DNA was extracted from mouse tissues using DNAeasy Blood and Tissue kit (Qiagen). The numbers of mice used in each individual experiment were calculated to permit detection of at least a two- to four-fold difference in bacterial loads between groups with 95% (two-sided) confidence and a power of 80%, based on prior experience.

### Ethics

Animal procedures were reviewed and approved by The Walter and Eliza Hall Institute of Medical Research Animal Ethics Committee (ethics approval number 2017.016) and were conducted in accordance with the Prevention of Cruelty to Animals Act (1986) and the Australian National Health and Medical Research Council Code of Practice for the Care and Use of Animals for Scientific Purposes (1997).

## Results

### General characterisation of *Mycobacterium spongiae* FSD4b-SM

We first performed general phenotypic analyses of *M. spongiae* FSD4b-SM that showed it was capable of growth on solid media typically used for culturing heterotrophic marine bacteria, although colony formation was scant (Fig. 1A). FSD4b-SM did not grow on media typically used for mycobacterial growth, such as Lowenstein-Jensen or egg-yolk-based agar media. An analysis of growth performed in simplified marine broth, showed that FSD4b-SM grew optimally at 28°C with an estimated doubling time of 64 days and reached stationary phase after approximately three months (Fig. 1B). FSD4b-SM was unable to grow at 37°C. The bacteria stained acid-fast, forming short, compact rods (Fig. 1C). Transmission electron microscopy confirmed rod-shaped cells, approx. 2 µM in length and 0.4 µM in diameter (Fig. 1D).

**Fig. 1:**
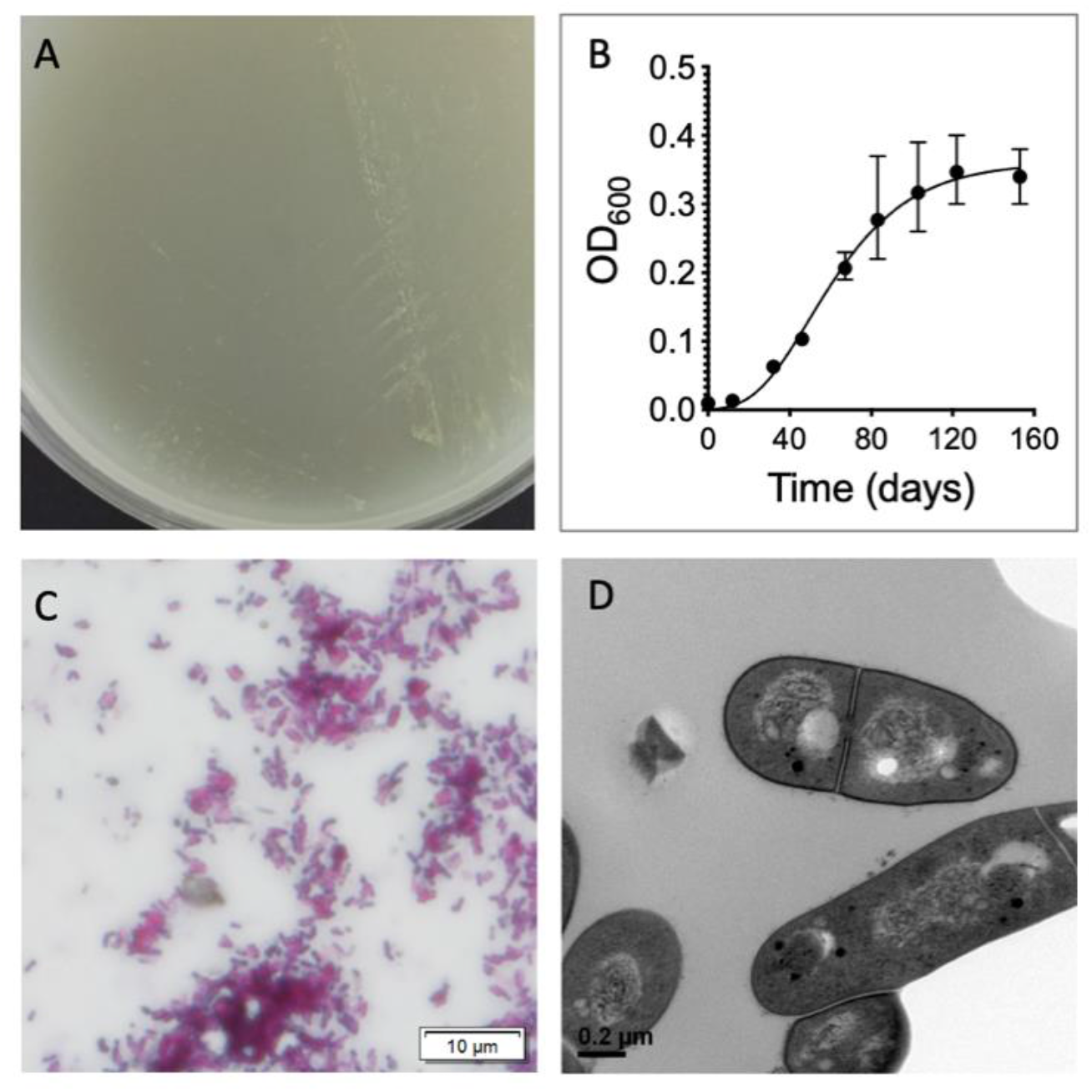
General phenotypic characteristics of *Mycobacterium* strain FSD4b-SM. A) Representative example of scant *M. spongiae* growth on simplified marine agar. B) Growth curve of *M. spongiae* in simplified marine broth. C) Ziehl-Neelsen stained *M. spongiae* cells. D) Electron micrograph of *M. spongiae* cells (x 33,000 magnification).

### Comparative genomics of *Mycobacterium spongiae* FSD4b-SM with other mycobacteria

Comparative genomics between related mycobacteria has provided significant insights into the factors that make *M. tuberculosis* pathogenic. Previous work using partial, concatenated 16S rRNA, *hsp65* and *rpoB* gene sequences from FSD4b-SM showed that it was closely related to the MTBC (8). To gain greater insight into the relationship between FSD4b-SM, the MTBC and other mycobacterial species, we first completed a high-quality, closed FSD4b-SM genome using a combination of Illumina and PacBio sequencing. FSD4b-SM has a single circular 5,581,157 bp chromosome, harbouring 4458 CDS (134 predicted pseudogenes) and a single rRNA locus. The average GC percentage was 65.56%. Initial comparisons of genome size showed that the FSD4b-SM genome is 1.1 Mb larger than the *M. tuberculosis* H37Rv genome (4.4Mb) (26, 27), but smaller than other close *M. tuberculosis* relatives, such as *Mycobacterium kansasii* (6.4 Mb) (28) and *Mycobacterium marinum* (6.6 Mb) (2).

To explore the relationship between FSD4b-SM and other mycobacteria more thoroughly, we calculated pairwise average nucleotide identity (ANI) with *M. tuberculosis* and eight other mycobacterial species known to be closely related to the MTBC (Fig. 2A). This analysis showed FSD4b-SM clustered most closely with *M. decipiens* and the MTBC (represented by *M. tuberculosis* and *M. canettii*) (approx. 80% ANI), however the overall ANI differences between all 10 mycobacterial genomes were minimal and cluster resolution was subsequently low (Fig. 2A). To assess the evolutionary relationship between FSD4b-SM, the MTBC and other mycobacteria we inferred a phylogeny among the same 10 mycobacteria and included an additional 20 comparator mycobacterial genomes (Fig. 2B) (4). A maximum-likelihood phylogeny built from an amino acid sequence alignment of 107 core CDS among all 30 mycobacterial genomes showed that FSD4b-SM clusters with *Mycobacterium decipiens* and the MTBC, within a group of mycobacteria previously defined as the *M. tuberculosis*-associated phylotype (MTBAP) and consistent with the ANI results (Fig. 2B) (4). The relatively long branch length of FSD4b-SM within the MTBAP cluster supports the classification of this mycobacterium as a distinct species (Fig. 2B). To delve further into the relationship between *M. tuberculosis* and FSD4b-SM, we more closely assessed core genome differences between these two mycobacteria and the genomes from three other key mycobacterial species (*M. kansasii, M. marinum, and M. decipiens*) (Fig. 2C). The five species shared a core genome of 1815 CDS, with FSD4b-SM sharing more than 50% of its CDS content with *M. marinum,* despite being more distantly related at the nucleotide level (Fig. 2C). However, *M. spongiae* and *M. tuberculosis* also share approximately 55% of their coding capacity; it is notable that a mammalian host-restricted pathogen like *M. tuberculosis* shares much of its protein coding capacity with a marine mycobacterium. Overall chromosome architecture is well conserved between FSD4b-SM and the closely related species, but there are also several regions spanning approximately 500kb in total of the FSD4b-SM genome that are distinct to the sponge mycobacterium (Fig. 2D, E). Upon further investigation it was noted that several of these distinct regions harboured large gene clusters encoding putative polyketide synthase (PKS) or non-ribosomal synthetase (NRPS) enzymes. These enzymes are involved in the production of specialised metabolites in bacteria and fungi, including many well-known bioactive molecules, such as antibiotics and anticancer compounds (29). To assess these PKS and NRPS regions further, we utilised the bioinformatic tool antiSMASH (17), which predicted a total of 19 regions potentially involved in specialised metabolism in the FSD4b-SM genome. As PKSs are heavily involved in the biosynthesis of core mycobacterial lipids, such as the mycoketides, phthiocerol dimycocerosates and mycolic acids (discussed in detail below), the majority of these 19 regions are well conserved across a range of mycobacteria. The FSD4b-SM genome also encodes an orthologous NRPS locus for the production of isonitrile lipopeptides (INLPs) (F6B93_01120 - 01155), which are used by *M. tuberculosis* and other pathogenic mycobacteria for metal transport (30). However, five of these 19 specialised metabolite loci appear to be specific to FSD4b-SM, corresponding to a total of 315 kb of DNA. These include a hybrid PKS-NRPS locus that has greater homology to PKS-NRPS systems from algae that those from other mycobacteria (F6B93_00330 – F6B93_00460); a putative alkylresorcinol locus (F6B93_10605 – F6B93_10760), that are known to produce molecules with antioxidant, cytotoxic or also have signalling properties (31); and three further PKS-encoding regions (F6B93_18505 – F6B93_18675, F6B93_19200 – F6B93_19380, F6B93_21085 – F6B93_21320) with unknown predicted products. At present it is not possible to predict the final molecular structures of a given PKS from genome sequence information alone, meaning that these regions await the identification of their ultimate chemical entities.

**Fig. 2:**
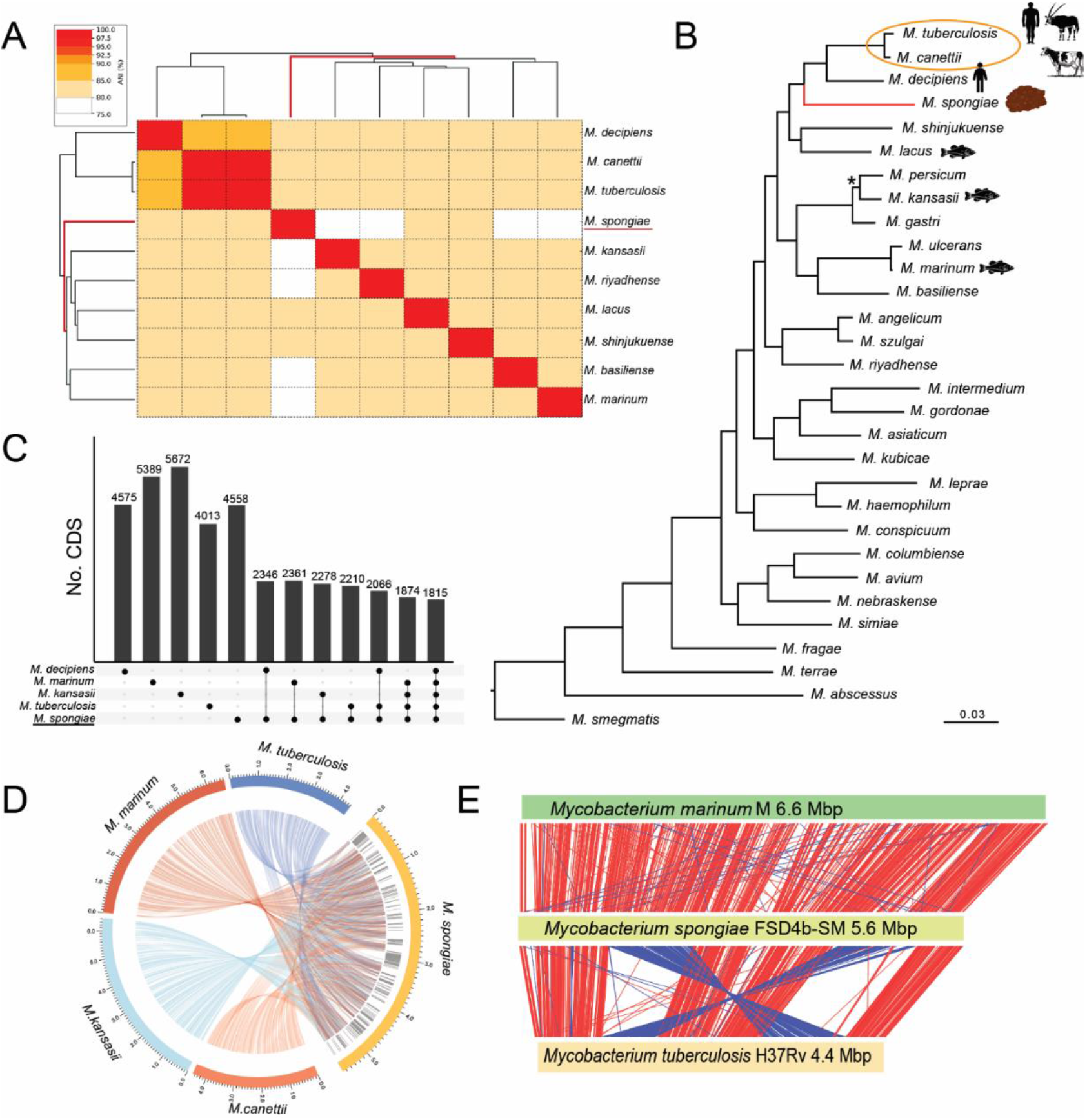
Comparative genomics summary of *Mycobacterium spongiae* FSD4b-SM. A) Pairwise average nucleotide identity (%ANI) between related *Mycobacterium* species. B) Maximum-likelihood phylogenetic tree (*iqtree*) with 1000 bootstrap iterations, inferred among 30 mycobacteria and based on amino acid sequence alignments from 107 conserved bacterial genes (*bcgTree*). *M. smegmatis* was used as an outgroup to root the phylogeny. Asterisk indicates node with >60% bootstrap node support. All other tree nodes had greater than 90% bootstrap support. MTBC encircled. Red branch length denotes *M. spongiae* FSD4b-SM placement. C) Upset plot showing shared coding sequences (CDS) at the 80% amino acid identity level between five related mycobacteria (*Roary*). (D) Circos plot summary of DNA sequence homology (*Blastn*) among *M. spongiae* and closely related species, showing regions (grey shading) ‘specific’ for *M. spongiae* among these comparisons. (E) ACT plot (Artemis) showing comparative chromosome architecture and length between *M. spongiae, M. marinum and M. tuberculosis*.

### Overview of shared *M. spongiae* and *M. tuberculosis* genetic features

Given the high level of genetic similarity and shared protein orthologues between *M. spongiae* and the MTBC, we sought to identify the presence of known *M. tuberculosis* pathogenesis factors in the FSD4b-SM genome. The *M. tuberculosis* antigens MPT83, TB8.4, antigen 85 complex, ESAT-6 and CFP-10, were all identified in the *M. spongiae* genome with >78% amino acid identity (Table S4). All four mammalian cell entry (Mce) families present in *M. tuberculosis* are conserved in gene content and synteny in FSD4b-SM, as well as a distinct *mce* locus not found in either *M. tuberculosis* or *M. marinum* (F6B93_18965 to F6B93_18990)(32) (Table S4). Other conserved pathogenic factors include the ESX secretion systems and a range of key mycobacterial lipid species (both discussed in detail below). Conserved regulatory systems include the PhoPR virulence regulatory system (33, 34) and several TetR-family regulators involved in *M. tuberculosis* antibiotic resistance such as EthR (and the associated monooxygenase EthA linked to ethionamide resistance), InhR (isoniazid) and EtbR (isoniazid and ethambutol) (35).

In *M. tuberculosis* the DosS-DosT/DosR (DevST/R) regulatory system controls approximately 50 CDS involved in responses to carbon monoxide (CO) and nitric oxide (NO) exposure (36). In FSD4b-SM a DosR ortholog exists (F6B93_06220), while F6B93_12710 is a putative sensor kinase most closely related to the *M. tuberulcosis* hypoxia sensor DosT (76% amino acid identity). A DosS orthologue appears absent in FSD4b-SM suggesting an inability to sense redox signals in the same way as *M. tuberculosis* (37). Other *M. tuberculosis* pathogenic determinants that are not well conserved in FSD4b-SM include a sphingomyelinase (encoded by Rv0888) that is used to degrade the major eukaryotic lipid sphingomyelin, and the *M. tuberculosis* outer-membrane channel protein and necrotizing exotoxin CpnT (Rv3903c), whose orthologue in FSD4b-SM (F6B93_13125) shares only 56% aa identity with only the NAD+-glycohydrolase domain of Rv3903c (38–40).

The MTBC has more than 80 toxin-antitoxin (TA) systems, important for bacterial persistence within the host. FSD4b-SM has a smaller and distinct toxin-antitoxin repertoire compared to the MTBC, with only 11 type II TA systems (F6B93_: 00510/00515, 00780/00785, 03975, 12335, 12350, 12355/12360, 13370/13375, 14995/15000, 15135, 18675/18680, 19810/19815). There are also no plasmids, phage or other insertion sequence elements in the FSD4b-SM genome in contrast to *M. tuberculosis* where mobile elements make up 3.4% of its genome (41). The absence of mobile DNA is also in contrast to the plasmids and prophage that appear in strains of *M. marinum* and *M. kansasii* (2, 3). This suggests that foreign DNA uptake has been restricted in FSD4b-SM. The FSD4b-SM genome contains a number of antibiotic resistance determinants (such as RbpA and aminoglycoside 2’-*N*-acetyltransferase) that are also present in *M. tuberculosis* (42, 43), inferring that these determinants are ancestral to this lineage of mycobacteria.

### DNA methylation in FSD4b

The absence of prophage, insertion sequence elements and plasmids suggest strong barriers in FSD4b-SM to extracellular DNA acquisition. DNA restriction modification is one possible barrier, so we took advantage of the PacBio sequence data to explore adenine DNA methylation patterns in this mycobacterium. We observed three different methylated motifs, two previously reported in *M. tuberculosis*, including the highly methylated CTCAG/CTGGAG motif (2582/2592 sites), and GTAYN4ATC (538/565 sites) (Table S5) (44). A third FSD4b-SM motif AGCN5CTTC/GAAGN5GCT (624/625 sites) is different to the third *M. tuberculosis* motif (Table S5). Interestingly, all three motifs had near complete methylation, suggesting efficient methylases working with their cognate (and presumably highly active) restriction modification systems.

### FSD4b-SM energetics

There is extensive conservation in FSD4b-SM with *M. tuberculosis* CDS encoding key proteins required for respiration and ATP synthesis, with CDS encoding all key proteins associated with the mycobacterial electron transport chain and the F_1_F_o_ ATP synthase present and intact (Fig. S1). However, like other mycobacteria outside the MTBC, a specific fumarate reductase complex (FrdABCD) is absent in *M. spongiae*. Also, the single nitrate reductase locus *nar* (F6B93_02320 - F6B93_02360) is distinct to that found in *M. tuberculosis*. The *M. tuberculosis hyc* hydrogenase locus encoding a purported formate hydrogenylase enzyme complex is also absent in FSD4b-SM (45). *M. spongiae* instead carries a locus encoding a group 1h hydrogenase complex (F6B93_RS07495 - F6B93_RS07595), orthologous to the *hhy* locus in *M. smegmatis* (46). This complex is presumably used by FSD4b-SM to oxidize molecular hydrogen that is abundant in seawater (47, 48).

### ESX systems

ESX (or type VII) secretion systems allow mycobacteria to export virulence determinants and other substrates across their specialized cell envelope. Most pathogenic mycobacteria contain up to five of these ESX systems, which are believed to have evolved by horizontal transfer and gene duplication events (49). The best studied of these systems is ESX-1, which is used by *M. tuberculosis* to permeabilize the phagolysosome and is also responsible for the processing and secretion of two key virulence determinants, CFP-10 (*esxB*) and ESAT-6 (*esxA*). All components of ESX-1 are highly conserved in FSD4b-SM, including CFP-10 and ESAT-6 orthologues (Fig. S2 and Table S6). Additionally, the FSD4b-SM ESX-1 locus contains a nine gene insertion that harbours novel PE and PPE encoding genes (Fig. S2). The four remaining ESX loci are all well conserved in the FSD4b-SM genome and include the components necessary for iron siderophore uptake (ESX-3, (50)) and for mycobacterial outer membrane permeability and nutrient uptake (ESX-5, (51)).

To investigate whether these ESX systems were functional in FSD4b-SM, we first performed secretion analysis followed by SDS-PAGE as well as Western blot analysis with an array of antibodies against *M. tuberculosis* proteins that are secreted by ESX-secretion systems or commonly used loading controls. While we were unable to detect any specific protein using SigA, GroEL, EsxN, EspA, EsxA (ESAT-6), EsxB (CFP-10) (data not shown), staining with an antibody against PE_PGRS proteins revealed that this group of proteins is highly expressed by FSD4b-SM (Fig. S3). High levels of the proteins were also detected in the culture filtrate. Compared to *M. tuberculosis* fractions the expression of PE_PGRS proteins appear at higher molecular weights, which is like *M. marinum*, another marine-associated mycobacterium (52, 53).

To investigate expressed proteins in more detail, we performed proteomic analysis of cell free culture supernatants and whole cell lysates. These analyses confidently identified a total of 1354 expressed proteins, including several proteins of the ESX-1, ESX-2 and ESX-5 secretion systems (Table S6). ESX-1 substrates and components detected included the major secreted *M. tuberculosis* antigen CFP-10 (F6B93_22245), the chaperone EspB (F6B93_22290), the PE protein chaperone EspG1 (F6B93_22210) and the ESX-1 secretion regulator EspI (F6B93_22255) (54–57). The ESX-5 secretion system has been shown to be essential to slow growing mycobacteria (51, 58) and several substrates of this system were detected, including a full-length version of the substrate EsxM (F6B93_11055), which has been suggested to promote dissemination in ancestral *M. tuberculosis* lineages (59). In addition, proteins constituting the building blocks of the ESX-5 secretion apparatus were also detected, including EccB5 (F6B93_11005), EccC5 (F6B93_11010), EccD5 (F6B93_11065) and EccE5 (F6B93_11075) (Table S6) (60). These results suggest a degree of conservation among type VII secretion systems and functionality between *M. tuberculosis* and the sponge mycobacterium.

### PE/PPE proteins

In addition to substrates encoded within each ESX locus, each ESX system also secretes its own array of PE/PPE substrates, so named for the highly conserved Proline-Glutamate and Proline-Proline-Glutamate motifs present in their N-termini, respectively (61). While PE/PPE proteins are found across both saprophytic and pathogenic mycobacteria, the latter generally harbour more of these proteins and in *M. tuberculosis* they are implicated in diverse phenotypes, including nutrient acquisition, and a range of pro- and anti-immunity responses (50, 61–63). Several *M. tuberculosis* PE/PPE proteins are essential for bacterial growth under a range of *in vitro* and *in vivo* conditions (64, 65). Examination of the FSD4b-SM genome identified 179 PE and 82 PPE genes (Table S7) corresponding to over 10% of its coding capacity, compared to 99 PE and 69 PPE genes in *M. tuberculosis* (7% of the genome) (26, 61). This repertoire includes multiple members of the PE_PGRS and PE_MPTR subfamilies that are restricted to members of the MTBC (61, 66) and is consistent with the expansion of PE/PPE family proteins in slow growing mycobacteria (16).

FSD4b-SM contains a number of orthologues to PE/PPE proteins reported to have important roles in *M. tuberculosis* growth including PPE4 (F6B93_02120) and its secretion partner PE5 (FB693_02115), that participate in mycobactin-mediated iron acquisition (50, 67, 68), PPE62 (F6B93_06010) involved in heme and hemoglobin utilisation (69), and PE19 (F6B93_11045) associated with stress resistance (Table S7) (70). Likewise, multiple *M. tuberculosis* PPEs that are involved in virulence have orthologues in *M. spongiae* including PPE25 (F6B93_11025), PPE26 (F6B93_11035), PPE27 (F6B93_04645), PPE68 (F6B93_22240) and PE35 (F6B93_11875) (71–73).

FSD4b-SM also possesses a number of PE/PPE proteins that have been positively linked to the secretion of type VII substrates, including PPE38 (F6B93_14895), which is essential for the secretion of PPE_MPTR and PE_PGRS proteins in the *M. tuberculosis* complex and *M. marinum* (74), and PE8 (F6B93_17700) and PPE15 (F6B93_17705) (51, 75). However, there are some potentially important differences too. For example, PE_PGRS47 (Rv2741), known to inhibit autophagy during *M. tuberculosis* infection, is absent from FSD4b-SM (76).

PE/PPE proteins that are involved in nutrient acquisition and stress resistance, such as PPE51 (F6B93_06205) and PE19 (F6B93_11045) are also present in FSD4b-SM with high levels of homology to their *M. tuberculosis* counterparts (70, 77). At least 29 of the putative FSD4b-SM PE/PPE proteins were confirmed to be expressed by proteomic analysis, including a LipY orthologue (F6B93_06345, LipY), which is involved in virulence in *M. tuberculosis* (78) (Table S7). Overall, a large expansion of PE/PPE proteins, in particular those of the PE_PGRS and PPE-MPTR families, is seen in *M. spongiae*.

Unlike in *M. tuberculosis*, there are no IS element insertions associated with PE/PPE genes in FSD4b-SM, suggesting that the expansion of PE/PPE repertoire in this strain, relative to *M. tuberculosis*, has most likely occurred by gene duplication and homologous recombination (79, 80). Overall, the pattern of FSD4b-SM PE/PPE proteins follows that in other mycobacteria, with multiple, high-identity paralogues. The roles these PE/PPE proteins play in FSD4b-SM physiology remain to be discovered, but a role in protein export across the highly hydrophobic mycobacterial cell envelope is one candidate function (81).

### Lipid biosynthesis pathways

The unique mycobacterial cell envelope is known to contain more than 50 classes of lipids and forms a barrier against many antibiotics and host immune defences. While mycolic acids make up approximately 30% of the outer layer of mycobacterial cells, other lipids such as the phthiocerol dimycocerosates (PDIMs) and phenolic glycolipids (PGLs) on the outer cell surface have important roles in pathogenicity and virulence (82). Due to the importance of lipids in mycobacterial physiology and virulence, we undertook a genomic and lipidomic evaluation of lipid production in FSD4b-SM.

Lipid species that are conserved among all mycobacteria include the mycolic acids and mycocerosates. Multiple enzymes are involved in mycolic acid biosynthesis in mycobacteria and highly conserved orthologues of each, including those involved in methoxy and hydroxy mycolic acid production, were identified in the FSD4b-SM genome (Table S8). Nearly all enzymes involved in mycolic acid biosynthesis were detected by our proteomic analysis, as were multiple mycolic acid species in our lipidomic analysis (Fig. 3, Table S8, S9). Likewise, the key enzyme in mycocerosate biosynthesis, Mas (F6B93_07120 in FSD4b-SM), was also detected in FSD4b-SM by proteomics. Interestingly, FSD4b-SM Mas has higher amino acid similarity to *M. marinum* Mas than to *M. tuberculosis* Mas, including the presence of a conserved tryptophan within the enoylreductase (ER) domain (83), which leads to the production of 2 *S* configured methyl branches on mycocerosic acids, which have only previously been seen in *M. marinum* and *M. ulcerans* (84) (Table S10).

**Fig. 3.**
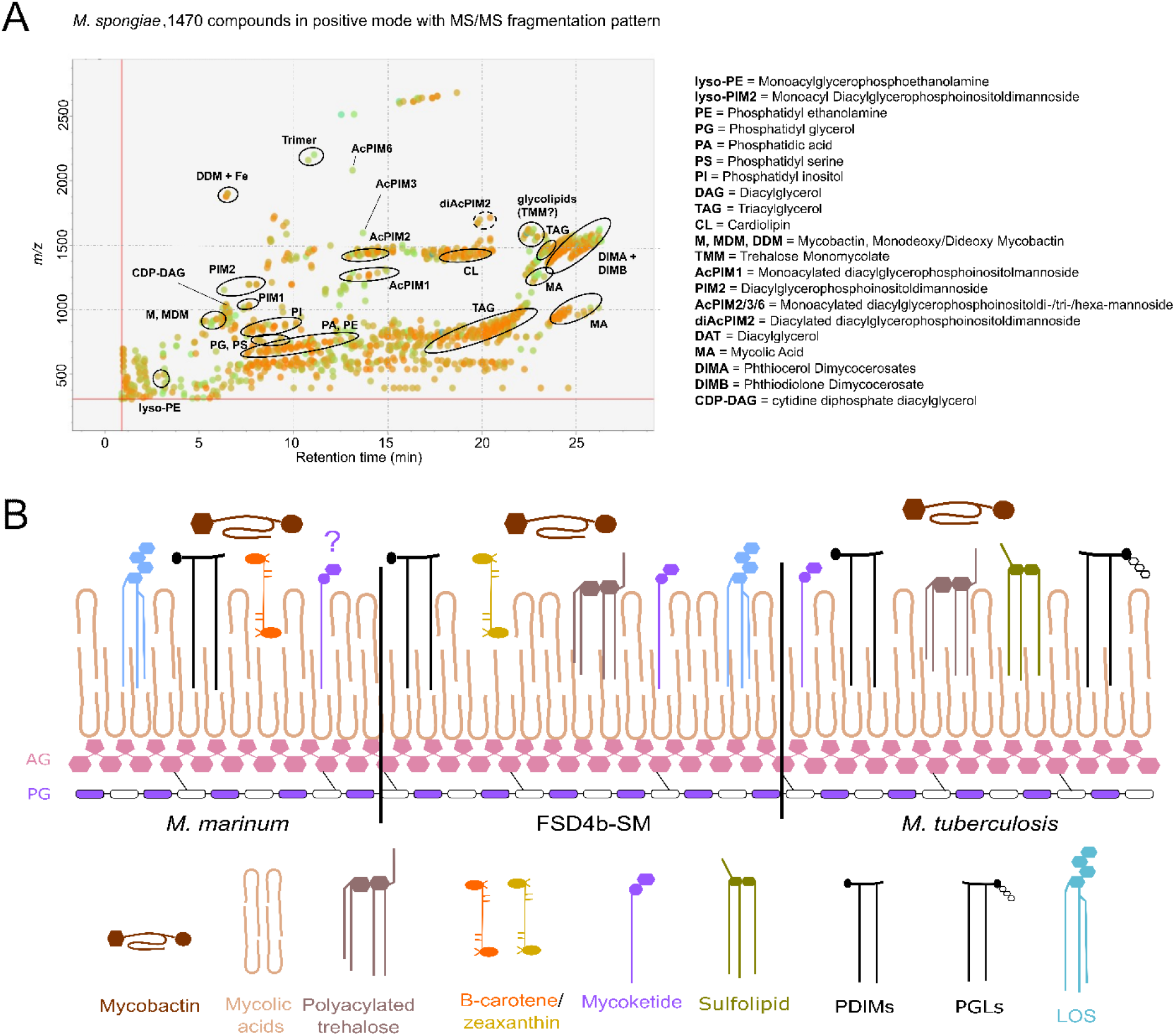
Lipidomic analysis of *M. spongiae*. A) MS/MS peak spot viewer of the *M. spongiae* extract in MS-DIAL showing retention time versus mass/charge ratio. Abundance (based on peak area) is shown as the colour of individual spots (blue = low, green = intermediate, orange = high abundance). Each lipid (sub)class is highlighted with a black circle. Note that not every spot within a circle belongs to the lipid (sub)class. B) Comparison of *M. marinum* and *M. tuberculosis* key membrane lipids with those predicted or detected from FSD4b-SM. The question mark above mycoketide for *M. marinum* indicates that the *M. marinum* genome contains a *pks12* orthologue, but mycoketides have not been detected from *M. marinum*. AG: arabinogalactan; PG: peptidoglycan; PDIMs: phthiocerol dimycocerosates; PGLs: phenolic glycolipids; LOS: lipooligosaccharide.

Genes encoding enzymes that produce lipid molecules restricted to pathogenic mycobacteria were also found in the FSD4b-SM genome. These include genes for mannosyl-β-1-phosphomycoketides (*pks12*), which are key lipid antigens presented by human CD1c T-cells during *M. tuberculosis* infection, and genes involved in the *M. tuberculosis*-restricted acylated trehalose derivatives including di-, tri- and poly-acyl trehalose (DAT, TAT and PAT, respectively), albeit with some potential structural variation (85–87). However, the putative DAT/TAT/PAT locus in FSD4b-SM is missing a critical FadD24 orthologue, which was found to be essential for both DAT and PAT formation in *M. tuberculosis* (88) (Fig. S4). DAT was not detected in our lipidomic analysis (Table S9), although this absence may also be due to possible differences in the putative DAT enzymes in FSD4b-SM and *M. tuberculosis* (see Table S11).

Genes for the production of the PDIMs were also identified, however, according to the biosynthetic logic (module and domain arrangement) of their producing polyketide synthases the FSD4b-SM enzymes should form unprecedented C9 D- and C11 L- configured phthiodiolone glycols in FSD4b-SM, in contrast to the L and D configuration seen in *M. tuberculosis* and the L and L configuration in *M. marinum* (Fig. S5, Table S10). However, these isomeric forms cannot be distinguished by our current lipidomic method. Genes for the formation of PGLs, which are thought to contribute to the hypervirulence of certain *M. tuberculosis* lineages are absent from FSD4b-SM and correspondingly no PGLs or their precursors were detected in our lipidomic analysis (Table S9, S10). Key genes for sulfolipid production appear to be only partially conserved in FSD4b-SM, suggesting that they are unlikely to be produced (Table S11).

Unlike pathogen-specific lipids, the highly antigenic lipooligosaccharides (LOS) have been detected from several slow growing mycobacterial saprophytes and pathogens, including *M. canettii* (38), but not in *M*. *tuberculosis sensu stricto* (89). Strains that make LOS contain a conserved genetic locus analogous to the DAT locus that contains two *pks* genes *(pks5* and *pks5.1*), FadD and Pap orthologues, as well as multiple glycosyltransferases (Table S11) (90–92). FSD4b-SM contains a LOS locus, which includes four *pks5* paralogues, compared to two in other species, suggesting a longer acyl chain may be added to the trehalose core (Fig. S7). In addition, FSD4b-SM also contains genes that appear to add a rhamnose to the acylated tetra-glucose core, suggesting that a previously unseen LOS is produced. FSD4b-SM also contains highly conserved genetic loci for the siderophore mycobactin and a *crt* locus, which is responsible for the production of the pigment beta-carotene in *M. marinum* (93) and *M. kansasii* (3). Lipidomic analysis detected the presence of the related isoprenoid pigment zeaxanthin in FSD4b-SM extracts (Table S8) and although zeaxanthin has not previously been detected from mycobacteria, it is known to be produced by many sea sponge-symbiotic bacteria (94, 95), where the pigment is believed to act as an antioxidant and protect sponges from UV-induced damage (95). These genomic, proteomic and metabolomic analyses of *M. spongiae* show that while its profile is distinct, it also has multiple features in common with *M. tuberculosis* as well as environmental opportunistic mycobacterial pathogens (Fig. 3).

#### Assessment of FSD4b-SM pathogenicity and its potential as a *M. tuberculosis* vaccine strain

At present, *M. bovis* BCG is the only licensed anti-tuberculosis vaccine that provides good protection against childhood forms of tuberculosis, but it is less efficient in protecting against pulmonary tuberculosis in adolescents and adults (96). A key reason for *M. bovis* BCG attenuation is the partial deletion of the ESX-1 locus and lack of major antigen ESAT-6 (97, 98). Early studies seeking to improve BCG by adding *M. tuberculosis* ESX-1 saw improved protection, but also increased virulence (98, 99). However, vaccine development using a rationally attenuated *M. tuberculosis* strain that still expresses ESAT-6 and CFP-10 showed increased protection relative to BCG, as did a recombinant BCG expressing the ESX-1 locus from the closely related *M. marinum* (57, 100). These studies are supported by a comprehensive analysis of recombinant BCG vaccines, which revealed that expression of ESX-1 derived effectors was an efficient way to improve BCG-induced immune responses (101). As the FSD4b-SM ESX-1 locus is more closely related to that from *M. tuberculosis* than *M. marinum*, we sought to investigate if *M. bovis* BCG carrying ESX-1^FSD4b-SM^, or alternatively the *M. spongiae* FSD4b-SM itself, would provide improved protection compared with BCG against *M. tuberculosis* aerosol challenge.

To construct a recombinant BCG strain expressing ESX-1^FSD4b-SM^ we used TAR cloning (102) to assemble the ESX-1 locus from eight overlapping PCR fragments in a yeast-*E. coli* shuttle vector and then subclone this region into the mycobacterial integrative vector pYUB412. Sequence analysis confirmed that the cloned region was identical to that from FSD4b-SM and this construct was integrated into *M. bovis* BCG at the phage L5 *attB* site. We then performed western blot analysis using ESAT-6 and CFP-10 specific antibodies. Only CFP-10 was detected in culture supernatants, a phenomenon observed with other recombinant BCG:ESX-1 expression systems (57). These experiments suggest that the ESX-1^FSD4b-SM^ was functional (Fig. 4).

**Fig. 4.**
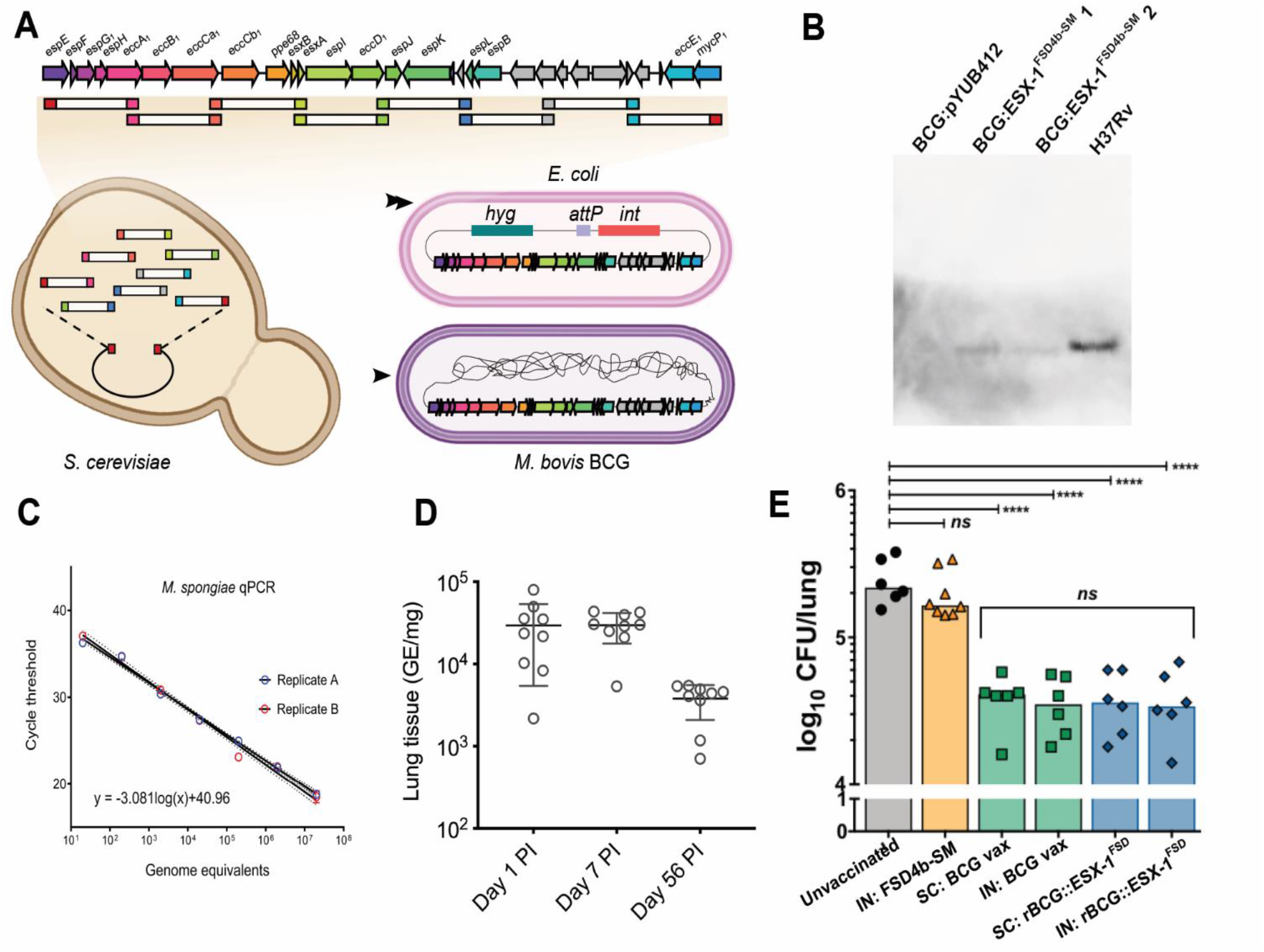
Construction of a recombinant *M. bovis* BCG with ESX-1FSD4b-SM and testing of its potential as an *M. tuberculosis* vaccine. A) The FSD4b-SM ESX-1 locus was assembled from eight overlapping PCR products in a yeast-*E.coli* shuttle vector in *Saccharomyces cerevisiae*, followed by transfer and sub-cloning of the ESX-1 locus into a mycobacterial integrative vector in *E. coli* and subsequent transfer and integration into the *M. bovis* BCG chromosome. B) Western blot with anti-CFP-10 antibody showing detection of CFP-10^FSD4b-SM^ from recombinant *M. bovis* BCG:ESX-1^FSD4b-SM^ and *M. tuberculosis* H37Rv CFP-10, but not from empty vector containing *M. bovis* BCG:pYUB412. BCG:ESX-1^FSD4b-SM^ 1 and 2 are two independent *M. bovis* BCG transformants. C) Establishment of a qPCR assay for detection of *M. spongiae* in mouse lung tissue. D) Mouse lung bacterial burden following intranasal inoculation of C57BL6 wild type mice with live *M. spongiae*. Shown are mean and SD at days 1, 7 and 56 post-infection (PI). E). Vaccination trial using *M. tuberculosis* H37Rv infectious aerosol challenge. Mice were vaccinated intranasally with *M. spongiae* FSD4b-SM or either intranasally or subcutaneously with *M. bovis* BCG (BCG vax), or recombinant *M. bovis* BCG expressing ESX-1 from *M. spongiae* (rBCG::ESX-1^FSD^).

With a CFP-10-expressing recombinant BCG in hand, we next sought to investigate whether this strain would improve protection against *M. tuberculosis* challenge in a murine infection model. We also sought to investigate whether live FSD4b-SM could be used as a potential vaccine candidate, as has been seen for other environmental mycobacteria (103). Due to the slow growth of FSD4b-SM, we developed a qPCR assay to monitor FSD4b-SM growth in mice. We performed both subcutaneous and intra-nasal vaccination routes. Interestingly, we observed *M. spongiae* FSD4b-SM persisted in the mouse lung for at least 8-weeks post inoculation (Fig. 4D). The mice also gained weight over that period and showed no signs of pathology.

Infectious challenge experiments with aerosolised *M. tuberculosis* H37Rv showed no protection offered by an intranasal dose of live *M. spongiae* (Fig. 4E). Recombinant *M. bovis* BCG:ESX-1^FSD4b-SM^ showed no further protection over the wild type *M. bovis* BCG vaccine, when delivered either subcutaneously or intranasally (Fig. 4E). The CD4^+^ T cell response against immunogenic regions of EsxA is a good correlate of protection against *M. tuberculosis* in C57BL/6 (H-2^b^) mice (99). We therefore explored the cross recognition of an EsxA:1-20 immunogenic region of *M. bovis* BCG:ESX-1^FSD4b-SM^ by a T cell hybridoma specific to *M. tuberculosis* EsxA, restricted by major histocompatibility complex molecule-II, and representative of CD4^+^ effector T cells induced in vivo. A T cell hybridoma specific to Ag85A was used as a positive infection control (Fig. S8A-C, top). Syngeneic DCs infected with recombinant *M. bovis* BCG:ESX-1^FSD4b-SM^ and co-cultured with NB11 T cell hybridomas (104) detected no cross recognition of EsxA from *M. bovis* BCG:ESX-1^FSD4b-SM^ (Fig. S8A-C, middle), in accordance with the poor conservation of the immunodominant EsxA:1-20 epitope in *M. spongiae* FSD4b-SM (Fig. S8D). However, as we also used an anti-EsxB T-cell hybridoma (XE12) that recognizes a 100% conserved EsxB region, restricted in the H-2^K^ haplotype (104), which was unable to detect *M. bovis* BCG:ESX-1^FSD4b-SM^-infected DCs (Fig. S8A-C, bottom), it is also possible that the EsxA and EsxB antigens from FSD4b-SM were not properly secreted by the recombinant BCG:ESX-1^FSD4b-SM^ strain under the infection conditions used in the experiment and thus not recognized. This may also explain why the use of this BCG:ESX-1^FSD4b-SM^ construct in vaccination experiments did not result in superior protection against *M. tuberculosis* challenge compared to BCG vaccination alone.

## Conclusion

*M. tuberculosis* and the MTBC have co-existed with humans for millennia. Our knowledge of the evolutionary trajectory that transformed an environmental mycobacterium into a host-adapted mammalian pathogen is enriched every time a new mycobacterium related to the MTBC is discovered (2–4, 105–108). Genomic reconstructions indicate *M. tuberculosis* evolved from a common ancestor shared with several aquatic environment-associated mycobacteria, including *M. marinum*, *M. kansasii* and *M. lacus* (2, 3). Here we have shown that a mycobacterium isolated from a marine sponge, for which we have proposed the name *Mycobacterium spongiae* sp. nov. (“of the sponge”), occupies a phylogenetic position even closer to *M. tuberculosis* than these other mycobacteria, thus adding further support to the hypothesis that the MTBC might have evolved from a marine mycobacterium. It is also interesting to consider that while sponges are not like humans, human lungs are somewhat like sponges at both a gross mechano-anatomical level (they are both biological filters) and also perhaps more profoundly at a molecular evolutionary level, as exemplified by the discovery of a conserved TNF-driven fibrinogenic response to silica exposure in sponges, present also in mammals where it can lead to silicosis (109). We don’t yet know anything of the interaction between *M. spongiae*, the host sponge from which it was isolated, *Fascaplysinopsis reticulata,* and its holobiont. Such interactions will be interesting to observe.

The close genetic relationship between FSD4b-SM and *M. tuberculosis* and functional similarities assessed by proteomics and lipidomics prompted us to examine whether FSD4b-SM or components thereof could provide protection against *M. tuberculosis* challenge in a murine lung-infection model. While previous studies have shown that *M. bovis* BCG expressing *M. marinum* ESX-1 provided superior protection to BCG alone in mice (100), this was not the case for the ESX-1 locus from FSD4b-SM, despite the fact that the key ESX-1 antigens ESAT-6 and CFP-10 from FSD4b-SM are more closely related to *M. tuberculosis* than those from *M. marinum*, although we do not know how well the *M. spongiae* ESX-1 locus is expressed in *M. bovis* BCG during *in vivo* growth conditions in the mouse. Moreover, looking at the genomic comparison map of the ESX-1 loci from different mycobacterial species, we noted that the *pe35* gene upstream of *ppe68* is not present in *M. spongiae* (Fig. S2). In *M. tuberculosis*, this gene commonly plays an important role in effector function, as *pe35* transposon mutants show ESAT-6 secretion defects and lower virulence (110, 111) although in such transposon mutants a possible effect of the transposon insertion on downstream effect might also be an explanation for this phenotype. However, the T cell hybridoma experiments were consistent with ESX-1 effector secretion defects and also help explain the poor vaccination outcome. Finally, the ESX-1 region of *M. spongiae* contains several extra genes compared to the ESX-1 loci of other mycobacteria (Fig. S2), whose potential impact on the functionally of the heterologous ESX-1 locus integrated into the BCG genome is unknown.

Overall, we describe a fascinating example of a slow growing, likely non-pathogenic mycobacterium we have designated *M. spongiae,* that is closely related to *M. tuberculosis*, sharing multiple genetic and functional similarities with the deadly human pathogen, while being adapted to the environmental conditions that prevail at 25 meters under the sea. It is presently unknown whether *M. spongiae* parasitises the sponge or only uses this ecological niche for its extracellular proliferation, but our findings strengthen the hypothesis that slow growing mycobacteria show an extraordinary capacity for adaptation to specific environments, a feature that has certainly also helped the ancestor of the tuberculosis-causing mycobacteria to adapt to mammalian hosts and conquer intracellular milieux.

### Description of *Mycobacterium spongiae* sp. nov

#### Mycobacterium spongiae (spon′gi.ae. L. gen. n. *spongiae* of a sponge, the source of the type strain)

Short, compact acid-fast staining rods approx. 2 µM in length and 0.4 µM in diameter. Capable of aerobic growth on solid media typically used for culturing heterotrophic marine bacteria, although colony formation is scant. Does not grow on media typically used for mycobacterial growth such as Lowenstein-Jensen or egg-yolk-based agar media. In a simplified marine broth (artificial seawater with 0.5% peptone, 0.1% yeast extract) grows optimally at 28°C with an estimated doubling time of 64 days and reaches stationary phase after approximately three months. Type strain is unable to grow at 37°C. Closely related to the MTBC (*Mycobacterium tuberculosis* complex) on the basis of 16S rRNA, *hsp65* and *rpoB* gene sequences. The type strain has a 16S rRNA gene similarity value of 99.6% with *Mycobacterium tuberculosis*. The ANI (pairwise average nucleotide identity) between reference genomes is supportive of the status of a species adjacent to the MTBC. The relatively long branch length of the type strain within the *M. tuberculosis*-associated phylotype (MTBAP) cluster by phylogenetic analysis based on amino acid sequence comparisons of 107 genes also supports distinct species status. Key *Mycobacterium tuberculosis* virulence factors are present, including intact ESX secretion systems and associated effectors. The genome size and number of coding DNA sequences of the type strain was 5,581,157 bp and 4458 genes (134 predicted pseudogenes). There is a single rRNA locus. The average G+C percentage based on the genome was 65.56%.

The type strain is FSD4b-SM.

## Acknowledgements

The authors wish to acknowledge the help of staff of the Mass Spectrometry facility at the Bio21 Institute, University of Melbourne. This project was supported by a National Health and Medical Research (NHMRC) L2 Fellowship to T.P.S., NHMRC Ideas grant (GNT2021638) to S.J.P and NHMRC Project Grant (GNT1105522) to T.P.S. and S.J.P. Research on sponge-associated bacteria in J.A.F.’s laboratory was funded by an Australian Research Council (ARC) Linkage project and the marine sponge collection was part of the Great Barrier Reef Seabed Biodiversity Project, research led at the Queensland Museum by John Hooper. H.I. was supported by a University of Queensland Research Scholarship (UQRS) and University of Queensland International Research Tuition Award (UQIRTA). The project also received support by the Agence Nationale pour la Recherche (ANR-10-LABX-62-IBEID) to RB.

