## Supplementary information for "A marine sponge-associated mycobacterium closely related to *Mycobacterium tuberculosis*"

**Contents:**

Table S1. Strains and plasmids used in this study.

Table S2. Genomes used for phylogenetic analysis.

Table S3. Primers used for construction of pYES1L:ESX-1$^{FSD4b-SM}$.

Table S4. *M. tuberculosis* virulence factors identified in *M. spongiae* genome

Table S5. Orthologues of *M. tuberculosis* ESX system proteins in *M. spongiae*

Figure S1. Analysis of energetics from *M. spongiae* genome

Figure S2. Alignment of ESX-1 and ESX-5 loci from *M. spongiae* and other related mycobacteria.

Table S6. Orthologues of *M. tuberculosis* ESX system proteins in *M. spongiae.*

Figure S3. Exuberant expression of PE_PGRS proteins in *M. spongiae*.

Table S7. PE/PPE proteins identified in FSD4b-SM and *M. tuberculosis* PE/PPE orthologues.

Table S8. Orthologues of mycolic acid biosynthesis proteins in *M. spongiae.*

Table S9**.** Summary of key lipid species identified in *M. spongiae*.

Table S10. Orthologues involved in the biosynthesis of phthiocerol dimycocerosates, phenolic glycolipids, and p-hydroxybenzoic acids.

Figure S4. Alignment of putative *pks3/4* locus in *M. spongiae* with that from *M. tuberculosis*.

Table S11. Orthologues of proteins involved in biosynthesis of trehalose and related compounds.

Figure S5. Predicted phthiodiolone diphthiocerante biosynthetic pathway in FSD4b-SM and comparison to phthiocerol dimycocerosate biosynthetic pathway in *M. tuberculosis*.

Figure S6. Alignment of *pks7-11* region in *M. tuberculosis* with that from *M. spongiae*.

Figure S7. Alignment of putative LOS locus in *M. spongiae* with those from other mycobacteria.

Figure S8. Assessment of recombinant *M. bovis* BCG FSD4b-SM Esx-1 protein effector T cell stimulation.

Table S12. Orthologues of proteins involved in the biosynthesis of phospholipids, isoprenoids and related compounds.

**Table S1**. Strains and plasmids used in this study.

| **Strain** | **Features** | Source |
|---|---|---|
| "*Mycobacterium spongiae sp. nov*" FSD4b-SM | | (1) |
| *M. bovis* BCG Danish (TMC 1010) | ATCC 35733 | ATCC |
| *Saccharomyces cerevisiae* MaV203 | *MATα; leu2-3,112; trp1-901; his3Δ200; ade2-101; cyh2R; can1R ; gal4Δ; gal80Δ; GAL1::lacZ; HIS3UASGAL1::HIS3@LYS2; SPAL10 UASGAL1::URA3.* | ThermoFisher Scientific |
| *Escherichia coli* TOP10 | F$^{-}$ *mcrA Δ(mrr-hsdRMS-mcrBC)* ϕ80*lacZΔM15 ΔlacX74 recA1 araD139 Δ(ara-leu) 7697 galU galK rpsL* (Str$^{R}$) *endA1 nupG* λ$^{-}$ | ThermoFisher Scientific |
| *Escherichia coli* DH10B | F$^{-}$ Δ(*ara-leu*) *7697*[Δ(*rapA'-cra'* )] Δ(*lac*)*X74*[Δ('*yahH-mhpE*)] duplication(514341-627601)[*nmpC-gltI*] *galK16 galE15* e14$^{-}$ (*icd*$^{WT}$ *mcrA*) ϕ80d *lacZΔM15 recA1 relA1 endA1* Tn*10.10 nupG rpsL150*(Str$^{R}$) *rph*$^{+}$ *spoT1 Δ(mrr-hsdRMS-mcrBC)* λ$^{-}$ | |
| **Plasmid** | **Features** | |
| pYES1L | *S. cerevisiae-E. coli* YAC-BAC shuttle vector. Contains ARS4/CEN5 ori for replication in yeast, TRP1 for selection in yeast and spectinomycin resistance for selection in *E. coli.* | ThermoFisher Scientific |
| pYES1L:ESX-1$^{FSD4b\text{-}SM}$ | Eight overlapping PCR fragments covering the ESX-1 locus from FSD4b-SM assembled into pYES1L. | This study |
| pYUB412 | E. coli-mycobacteria shuttle vector. Hyg$^{R}$ in mycobacteria, Amp$^{R}$ for selection in *E. coli*, ϕL5 *attP* site and integrase for integration into L5 *attB* site in mycobacterial chromosomes. | (2) |
| pYUB412:ESX-1$^{FSD4b\text{-}SM}$ | ESX-1$^{FSD4b\text{-}SM}$ cloned from pYES1L:ESX-1$^{FSD4b\text{-}SM}$ by *Sbf*I digestion into *Sbf*I site in pYUB412. | This study |

**Table S2**. Genomes used for phylogenetic analysis.

| **Species** | **Strain** | **Accession number** |
|---|---|---|
| *M. angelicum* | DSM 45057 | GCF_002086155.1 |
| *M. asiaticum* | DSM 44297 | GCF_000613245.1 |
| *M. avium* | 104 | GCF_000014985.1 |
| *M. basilense* | 901379 | GCF_900292015.1 |
| *M. canettii* | STB-K | GCF_000328785.1 |
| *M. colombiense* | CECT 3035 | GCF_002105755.1 |
| *M. conspicuum* | DSM 44136 | GCF_002102095.1 |
| *M. decipiens* | TBL 1200985 | GCF_002104675.1 |
| *M. fragae* | DSM 45731 | GCF_002102185.1 |
| *M. gastri* | DSM 43505 | GCF_002102175.1 |
| *M. gordonae* | DSM 44160 | GCF_002101675.1 |
| *M. haemophilum* | DSM 44634 | GCF_000340435.2 |
| *M. intermedium* | DSM 44049 | GCF_002086275.1 |
| *M. kansasii* | ATCC 12478 | GCF_000157895.3 |
| *M. kubicae* | CIP 106428 | GCF_002101745.1 |
| *M. lacus* | DSM 44577 | GCF_002102215.1 |
| *M. leprae* | TN | GCF_000195855.1 |

| | | |
|---|---|---|
| *M. marinum* | M | NC_010612 |
| *M. nebraskense* | DSM 44803 | GCF_002102255.1 |
| *M. persicum* | AFPC-000227 | GCF_002086675.1 |
| *M. riyadhense* | DSM 45176 | GCF_002101845.1 |
| *M. shinjukuense* | CCUG 53584 | GCF_002086755.1 |
| *M. simiae* | MO323 | GCF_001584765.1 |
| *M. smegmatis* | MC2 155 | GCF_000015005.1 |
| *M. szulgai* | DSM 44166 | GCF_002116635.1 |
| *M. terrae* | CIP 104321 | GCF_002101955.1 |
| *M. tuberculosis* | H37Rv | GCF_000195955.2 |
| *M. ulcerans* | Agy99 | CP000325 |

**Table S3**. Primers used for construction of pYES1L:ESX-1$^{\text{FSD4b-SM}}$.

| Oligo name | Sequence (5’ → 3’) | Comment |
|---|---|---|
| FSD4b ESX-1 R1-F | CGCAGCGGCGGCCGCGCTGATACCGCCGCCCTGCAGGACTA GTTCGGTTTTAGAACGTG | PCR ESX-1, region 1 for yeast assembly. Overlaps with vector pYES1L. |
| FSD4b ESX-1 R1-R | GCAGACCGCGCCTTTTGCCGTGCGAGATCGGCAATCACG | PCR ESX-1, region 1 for yeast assembly. Overlaps with 5’ end of region 2 product. |
| FSD4b ESX-1 R2-F | CGATCTCGCACGGCAAAAGGCGCGGTCTGCCCAATACAC | PCR ESX-1, region 2 for yeast assembly. Overlaps with 3’ end of region 1 product. |
| FSD4b ESX-1 R2-R | GGTGAGTACCCGGATCTCCAGTAGGTCGGGCGGATGAAA | PCR ESX-1, region 2 for yeast assembly. Overlaps with 5’ end of region 3 product. |
| FSD4b ESX-1 R3-F | CCCGACCTACTGGAGATCCGGGTACTCACCGAGAACCCC | PCR ESX-1, region 3 for yeast assembly. Overlaps with 3’ end of region 2 product. |
| FSD4b ESX-1 R3-R | CTCCGGCAGCGACGGTGTCGTTGCCATCGCCTGCGAGTA | PCR ESX-1, region 3 for yeast assembly. Overlaps with 5’ end of region 4 product. |
| FSD4b ESX-1 R4-F | GCGATGGCAACGACACCGTCGCTGCCGGAGATCGCCGCT | PCR ESX-1, region 3 for yeast assembly. Overlaps with 3’ end of region 3 product. |
| FSD4b ESX-1 R4-R | CGACCGCGTCGATGACGTCCTCGACGAGTGGCCGATAGC | PCR ESX-1, region 3 for yeast assembly. Overlaps with 5’ end of region 5 product. |
| FSD4b ESX-1 R5-F | CACTCGTCGAGGACGTCATCGACGCGGTCGCAGTACTCG | PCR ESX-1, region 3 for yeast assembly. Overlaps with 3’ end of region 4 product. |
| FSD4b ESX-1 R5-R | AGGCGGTATTTGGTCCGGTGGGGCGGCCAGTGCCGCCAA | PCR ESX-1, region 3 for yeast assembly. Overlaps with 5’ end of region 6 product. |
| FSD4b ESX-1 R6-F | CTGGCCGCCCCACCGGACCAAATACCGCCTTCGAAGACT | PCR ESX-1, region 3 for yeast assembly. Overlaps with 3’ end of region 5 product. |
| FSD4b ESX-1 R6-R | TACTAGGCCTTGGTGTGGGCAGTTCTGCTGCCGCGGAGG | PCR ESX-1, region 3 for yeast assembly. Overlaps with 5’ end of region 7 product. |
| FSD4b ESX-1 R7-F | CAGCAGAACTGCCCACACCAAGGCCTAGTACATCAGCTT | PCR ESX-1, region 3 for yeast assembly. Overlaps with 3’ end of region 6 product. |
| FSD4b ESX-1 R7-R | GGTAGATCGTGTCTGAAGGTGTTCCACCGTAGAACGTCA | PCR ESX-1, region 3 for yeast assembly. Overlaps with 5’ end of region 8 product. |
| FSD4b ESX-1 R8-F | ACGGTGGAACACCTTCAGACACGATCTACCCGACGGCCA | PCR ESX-1, region 3 for yeast assembly. Overlaps with 3’ end of region 7 product. |
| FSD4b ESX-1 R8-R | TCACTGACTTTAATTAACTGCGGCGAGGCCTGCAGGCACCGC GGTTGTCGCCGGGCTGG | PCR ESX-1, region 3 for yeast assembly. Overlaps with vector pYES1L. |

**Table S4**. *M. tuberculosis* virulence factors identified in *M. spongiae* genome

| H37Rv locus tag | Protein | Function | *M. spongiae* locus tag | % identity | Identified by proteomics |
|---|---|---|---|---|---|
| Rv2873 | MPT83 | Unknown | F6B93_07425 | 80 | |
| Rv1174c | TB8.4 | Malonyl-CoA:AcpM transacylase | F6B93_17085 | 83 | |
| Rv3804c | FbpA (Ag85A) | Mycolyltransferase, TDM/mAGP assembly | F6B93_21945 | 90 | Y |
| Rv1886c | FbpB (Ag85B) | Mycolyltransferase, TDM/mAGP assembly | F6B93_11510 | 89 | Y |
| Rv0129c | FbpC (Ag85C) | Mycolyltransferase, TDM/mAGP assembly | F6B93_01240 | 86 | Y |
| Rv3874 | CFP10 (EsxB) | | F6B93_22245 | 84 | Y |
| Rv3875 | ESAT-6 (EsxA) | | F6B93_22250 | 80 | |
| Rv0757 | PhoP | Transcriptional regulator | FSD4b_04125 | 95 | |
| Rv0758 | PhoR | Sensor kinase | FSD4b_04124 | 77 | |
| Rv0169 | Mce1A | Mce-family protein | F6B93_01525 | 83 | Y |
| Rv0170 | Mce1B | Mce-family protein | F6B93_01530 | 88 | Y |
| Rv0171 | Mce1C | Mce-family protein | F6B93_01535 | 89 | Y |
| Rv0172 | Mce1D | Mce-family protein | F6B93_01540 | 89 | Y |
| Rv0173 | LprK | MCE-family lipoprotein | F6B93_01545 | 84 | Y |
| Rv0174 | Mce1F | Mce-family protein | F6B93_01550 | 89 | Y |
| Rv0589 | Mce2A | Mce-family protein | F6B93_03845 | 84 | |
| Rv0590 | Mce2B | Mce-family protein | F6B93_03850 | 85 | |
| Rv0591 | Mce2C | Mce-family protein | F6B93_03855 | 78 | |
| Rv0592 | Mce2D | Mce-family protein | F6B93_03860 | 83 | |
| Rv0593 | LprL | MCE-family lipoprotein | F6B93_03865 | 81 | |
| Rv0594 | Mce2F | Mce-family protein | F6B93_03870 | 84 | Y |
| Rv1966 | Mce3A | Mce-family protein | F6B93_11815 | 81 | |
| Rv1967 | Mce3B | Mce-family protein | F6B93_11820 | 85 | |
| Rv1968 | Mce3C | Mce-family protein | F6B93_11825 | 84 | |
| Rv1969 | Mce3D | Mce-family protein | F6B93_11830 | 83 | |
| Rv1970 | LprM | MCE-family lipoprotein | F6B93_11835 | 87 | |
| Rv1971 | Mce3F | Mce-family protein | F6B93_11840 | 80 | |
| Rv3499c | Mce4A | Mce-family protein | F6B93_20140 | 84 | Y |
| Rv3498c | Mce4B | Mce-family protein | F6B93_20135 | 84 | Y |
| Rv3497v | Mce4C | Mce-family protein | F6B93_20130 | 84 | Y |
| Rv3496c | Mce4D | Mce-family protein | F6B93_20125 | 84 | Y |
| Rv3495c | LprN | MCE-family lipoprotein | F6B93_20120 | 84 | Y |
| Rv3494c | Mce4F | Mce-family protein | F6B93_20115 | 82 | Y |

**Table S5**. DNA methylation patterns identified in the *M. spongiae* genome.

| Methylation type | # detected | # in genome | Sequence |
|---|---|---|---|
| m6A | 624 | 625 | AGCNNNNNCTTC/GAAGNNNNNGCT |
| m6A | 2582 | 2592 | CTCCAG/CTGGAG |
| m6A | 538 | 565 | GTAYNNNNATC/GATNNNNRTAC |

| | Operon/ Gene[1] | *Subunits* | Rv # | Enzyme Name |
|---|---|---|---|---|
| 86% | *ace* | - | Rv2241 | Pyruvate dehydrogenase E1 component |
| 83% | *lpd* | - | Rv0462 | Dihydrolipoamide dehydrogenase |
| | *hyc* | Rv0081<br>Rv0082<br>Rv0083<br>*hycDPQE*<br>Rv0088 Truncated | Rv0081-Rv0088 | Formate hydrogenlyase OR Energy-converting hydrogenase related complex (Ehr) (a.k.a HydTB) |
| 85% | *qcr* | *qcrC*<br>*qcrA*<br>*qcrB* | Rv2194<br>Rv2195<br>Rv2196 | Cytochrome $bc_1$ |
| 83% | *ctaB* | - | Rv1451 | Cytochrome c oxidase assembly factor |
| 81% | *ctaC* | *ctaC*<br>- | Rv2200c<br>*(Rv2199c)* | Transmembrane Cytochrome C subunit II |
| 86% | *ctaD* | - | Rv3043c | Cytochrome C oxidase polypeptide I |
| 87% | *ctaE* | - | Rv2193 | Cytochrome C oxidase subunit III |
| 80% | *cyd* | *cydA*<br>*cydB*<br>*cydD*<br>*cydC* | Rv1623c<br>Rv1622c<br>Rv1621c<br>Rv1620c | Cytochrome *bd* oxidase |
| 65% | *nar* | *narG*<br>*narH*<br>*narJ*<br>*narI* | Rv1161<br>Rv1162<br>Rv1163<br>Rv1164 | Menaquinone:Nitrate reductase |
| 83% | *sirA* | *sirA*<br>*cysH*<br>*cheI* | Rv2391<br>Rv2392<br>Rv2393 | Sulfite reductase<br>APS reductase<br>ferrochelatase |
| 81% | *nirBD* | *nirB*<br>*nirD* | Rv0252<br>Rv0253 | Nitrite reductase [NAD(P)H] |
| | *frd* | *frdA*<br>*frdB*<br>*frdC*<br>*frdD* | Rv1552<br>Rv1553<br>Rv1554<br>Rv1555 | Menaquinone:Fumarate reductase |
| 84% | *atp* | *atpIBEFHAGDC* and Rv1312 | Rv1303-Rv1312 | $F_1F_o$ ATP synthase operon |
| 83% | *nuo* | *nuoABCDEFGHIJKLMN* | Rv3145–Rv3158 | Type I NADH:Menaquinone oxidoreductase |
| 81% | *ndh* | - | Rv1854c | Type II NADH:Menaquinone oxidoreductase |
| 81% | *ndhA* | - | Rv0392c | Type II NADH:Menaquinone oxidoreductase |
| 83% | *sdh1* | *sdh1CD*<br>*sdh1A*<br>*sdh1B* | Rv0249c<br>Rv0248c<br>Rv0247c | Succinate:Menaquinone oxidoreductase I |
| 87% | *sdh2* | *sdh2C*<br>*sdh2D*<br>*sdh2A*<br>*sdh2B* | Rv3316<br>Rv3317<br>Rv3318<br>Rv3319 | Succinate:Menaquinone oxidoreductase II |
| 79% | *mqo* | - | Rv2852c | Malate:Menaquinone oxidoreductase |
| 81% | *pru* | *pruA*<br>*pruB* | Rv1187<br>Rv1188 | Proline dehydrogenase and pyrroline-5-carboxylate dehydrogenase |
| 84% | *gpdA1* | - | Rv0564c | Glycerol-3-phosphase dehydrogenase A1 |
| 82% | *gpdA2* | - | Rv2982c | Glycerol-3-phosphase dehydrogenase A2 |
| 79% | *glpD1* | -<br>- | Rv2249c<br>(Rv2250c) | Glycerol-3-phosphase dehydrogenase D1 |
| 83% | *glpD2* | *glpD2*<br>*ldpA* | Rv3302c<br>Rv3303c | Glycerol-3-phosphase dehydrogenase D2<br>Dihydrolipoamide dehydrogenase |
| 83% | *cox* | *coxCMSLDEFG* | Rv0376c–Rv0368c | Carbon monoxide dehydrogenase |
| 82% | *ald* | - | Rv2780 | L-Alanine dehydrogenase |
| 80% | *lldD1* | *?*<br>*pqqE*<br>*lldD1* | Rv0692<br>Rv0693<br>Rv0694 | L-Lactate dehydrogenase 1 |
| | *lldD2* | *lldD2* 80%<br>-<br>- Endonuclease | Rv1872c<br>Rv1871c<br>Rv1870c | L-Lactate dehydrogenase 2 |
| | *Rv0843* | - | (Rv0842)<br>Rv0843 | Pyruvate dehydrogenase E1 component alpha subunit |
| 80% | *pdh* | *pdhA*<br>*pdhB*<br>*pdhC* | Rv2497c<br>Rv2496c<br>Rv2495c | Pyruvate dehydrogenase |

**Figure S1**. Analysis of energetics from *M. spongiae* genome. Blue bars indicate conservation of a particular system with other mycobacteria, while red bars indicate absence of a particular system.

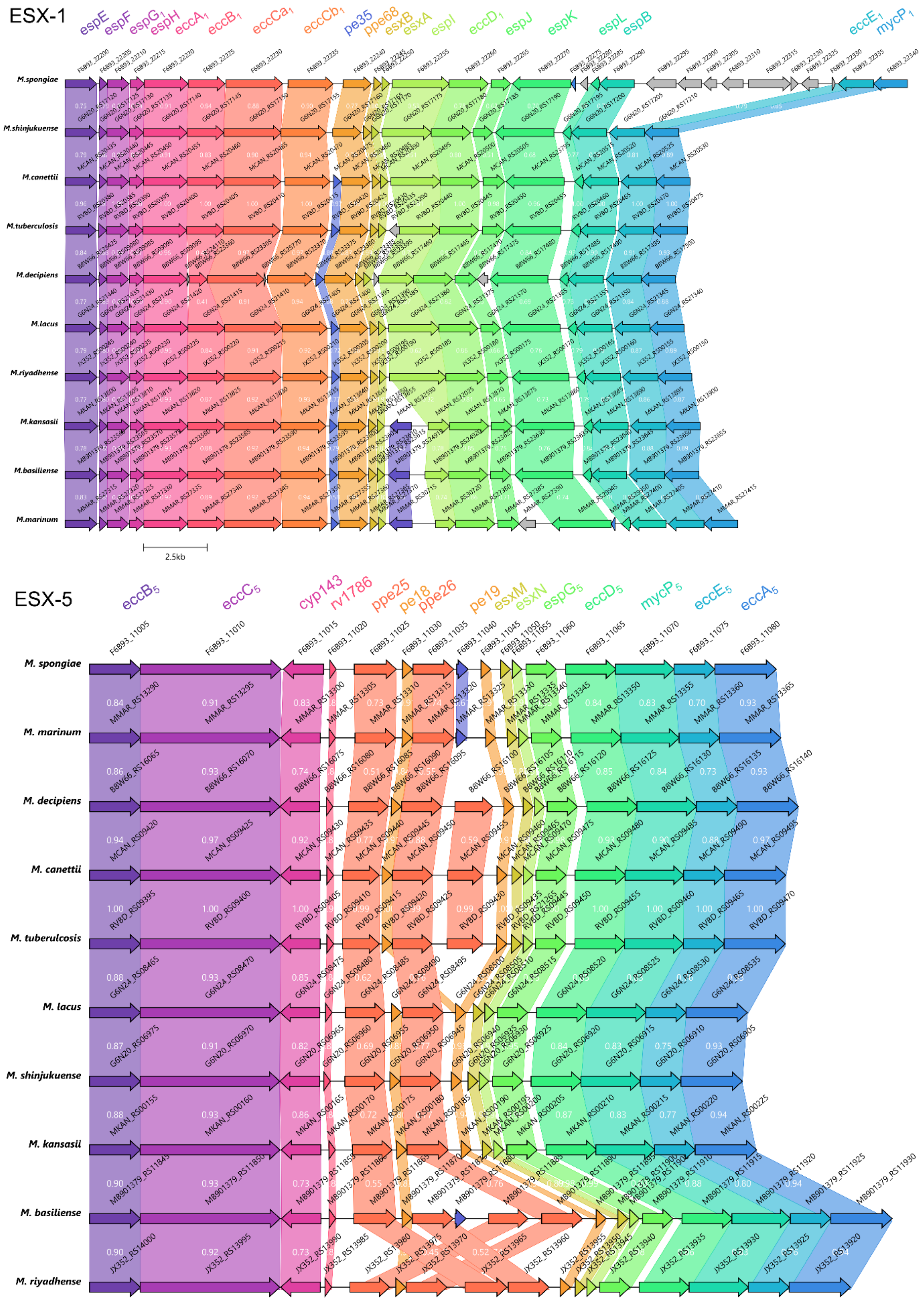

**Figure S2**. Alignment of ESX-1 and ESX-5 loci from *M. spongiae* FD4b-SM and other related mycobacteria. Pairwise percentage identity between encoded proteins is shown in white text, locus tags are written in black. Common gene names for each the Esx proteins are coloured according to the gene on which they are encoded.

**Table S6.** Orthologues of *M. tuberculosis* ESX system proteins in *M. spongiae*

| H37Rv locus orthologue | Description | *M. spongiae* locus tag | % identity | Detected by proteomics? |
|---|---|---|---|---|
| **ESX-1** | | | | |
| RV3864 | EspE | F6B93_22200 | 76 | |
| RV3865 | EspF | F6B93_22205 | 82 | |
| RV3866 | EspG1 | F6B93_22210 | 80 | Y |
| RV3867 | EspH | F6B93_22215 | 75 | |
| RV3868 | EccA1 | F6B93_22220 | 89 | |
| RV3869 | EccB1 | F6B93_22225 | 82 | |
| RV3870 | EccCa1 | F6B93_22230 | 90 | |
| RV3871 | EccCb1 | F6B93_22235 | 91 | |
| RV3872 | PE35 | F6B93_11875 | 57 | |
| RV3873 | PPE68 | F6B93_22240 | 72 | |
| RV3874 | CFP10 (EsxB) | F6B93_22245 | 84 | Y |
| RV3875 | ESAT-6 (EsxA) | F6B93_22250 | 80 | |
| RV3876 | EspI | F6B93_22255 | 69 | Y |
| RV3877 | EccD1 | F6B93_22260 | 78 | |
| RV3878 | EspJ | F6B93_22265 | 59 | |
| RV3879 | EspK | F6B93_22270 | 72 | |
| RV3880 | EspL | F6B93_22285 | 75 | |
| RV3881 | EspB | F6B93_22290 | 71 | Y |
| RV3882 | EccE1 | F6B93_22335 | 82 | |
| RV3883 | MycP1 | F6B93_22340 | 86 | |
| **ESX-2** | | | | |
| RV3884c | EccA2 | F6B93_22345 | 89 | |
| RV3885c | EccE2 | F6B93_22350 | 81 | |
| RV3886c | MycP2 | F6B93_22355 | 81 | |
| RV3887c | EccD2 | F6B93_22360 | 86 | |
| RV3888c | | F6B93_22365 | 90 | Y |
| RV3889c | EspG2 | F6B93_22370 | 89 | |
| RV3890c | EsxC | F6B93_22375 | 86 | |
| RV3891c | EsxD | F6B93_22380 | 90 | |
| RV3892c | PPE69 | F6B93_22385 | 81 | |
| RV3893c | PE36 | F6B93_22390 | 86 | |
| RV3895c | | F6B93_22395 | 36 | |
| RV2209 | | F6B93_22400 | 36 | |
| RV3894c | EccC2 | F6B93_22405 | 88 | |
| RV3895c | EccB2 | F6B93_22410 | 86 | |
| **ESX-3** | | | | |
| RV0282 | EccA3 | F6B93_02100 | 87 | |
| RV0283 | EccB3 | F6B93_02105 | 75 | |
| RV0284 | EccC3 | F6B93_02110 | 84 | |
| RV0285 | PE5 | F6B93_02115 | 88 | |
| RV0286 | PPE4 | F6B93_02120 | 67 | |
| RV0287 | EsxG | F6B93_02125 | 91 | |
| RV0288 | EsxH | F6B93_02130 | 85 | |
| RV0289 | EspG3 | F6B93_02135 | 80 | |
| RV0290 | EccD3 | F6B93_02140 | 82 | |
| RV0291 | MycP3 | F6B93_02145 | 70 | |
| RV0292 | EccE3 | F6B93_02150 | 65 | |
| **ESX-4** | | | | |
| RV3450c | EccB4 | FSD4b_00956 | 68 | |
| RV3449 | MycP4 | FSD4b_00957 | 69 | |
| RV3448 | EccD4 | FSD4b_00958 | 59 | |
| RV3447c | EccC4 | FSD4b_00959 | 65 | |
| RV3446c | | FSD4b_00960 | 55 | |
| RV3445c | EsxU | FSD4b_00961 | 78 | |
| RV3444c | EsxT | FSD4b_00962 | 80 | |
| **ESX-5** | | | | |
| RV1782 | EccB5 | F6B93_11005 | 87 | Y |
| RV1783 | EccC5 | F6B93_11010 | 91 | Y |

| | | | | |
|---|---|---|---|---|
| RV1785 | Cyp143 | F6B93_11015 | 74 | |
| RV1786 | | F6B93_11020 | 79 | |
| RV1787 | PPE25 | F6B93_11025 | 49 | |
| RV1788 | PE18 | F6B93_11030 | 86 | |
| RV1789 | PPE26 | F6B93_11035 | 53 | |
| RV1271 | | F6B93_11040 | 68 | |
| RV1791 | PE19 | F6B93_11045 | 95 | |
| RV1792 | EsxM | F6B93_11050 | 93 | Y |
| RV1793 | EsxN | F6B93_11055 | 94 | Y |
| RV1794 | | F6B93_11060 | 94 | Y |
| RV1795 | EccD5 | F6B93_11065 | 85 | Y |
| RV1796 | MycP5 | F6B93_11070 | 84 | |
| RV1797 | EccE5 | F6B93_11075 | 75 | Y |
| RV1798 | EccA5 | F6B93_11080 | 95 | Y |

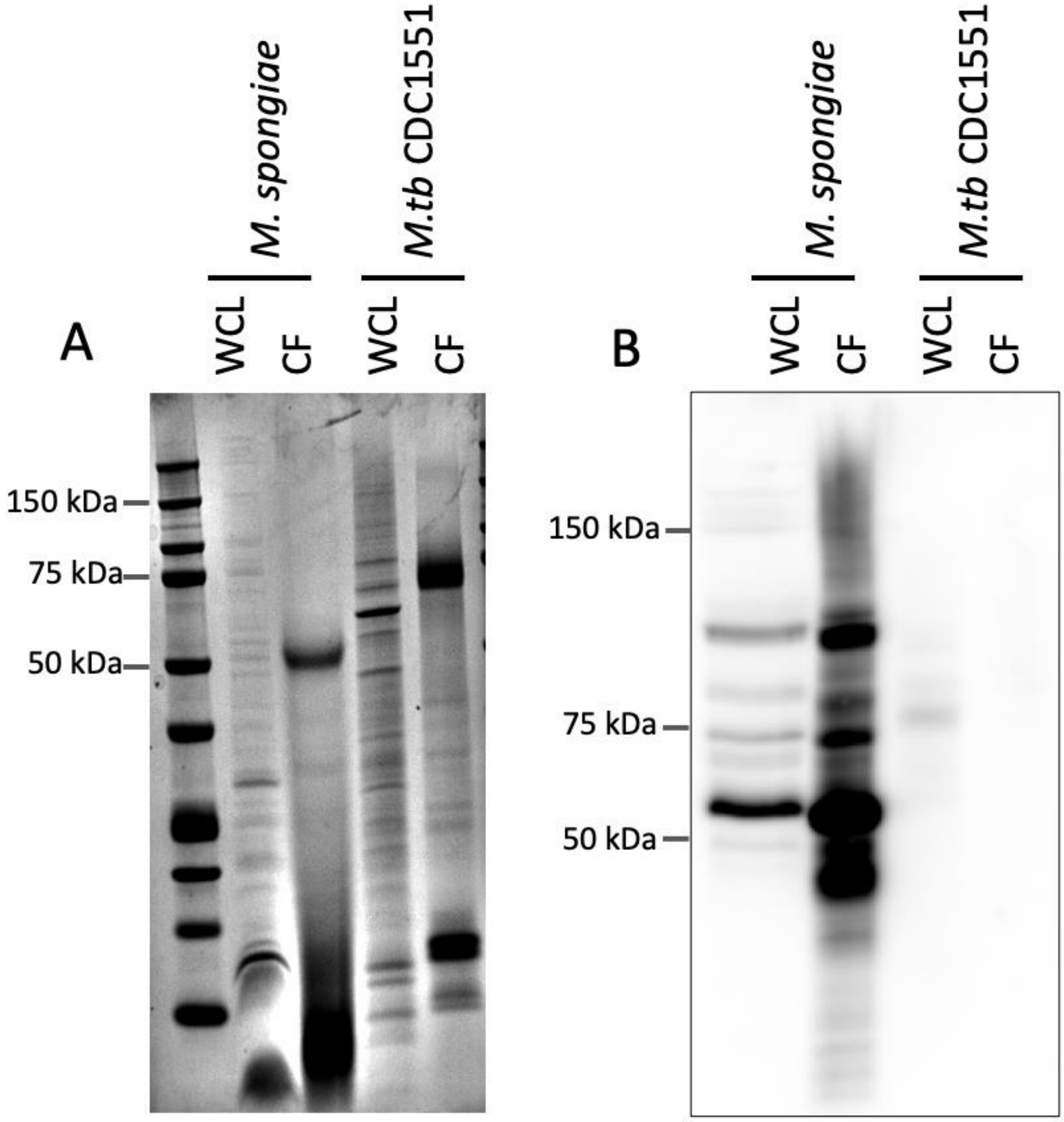

**Figure S3.** Exuberant expression of PE_PGRS proteins in *M. spongiae*. (A) SDS-PAGE, Coomassie brilliant blue stained gel, showing protein loading for *M. spongiae* and *M. tuberculosis*. (B). Western blot analysis using anti-PE_PGRS antibody against *M. spongiae* and *M. tuberculosis*. WCL = Whole Cell Lysate. CF = Culture Filtrate.

**Table S7. PE/PPE proteins identified in FSD4b-SM and *M. tuberculosis* PE/PPE orthologues**

Note: Proteins were selected for analysis via BLAST matches with known PE/PPE proteins or via annotation describing the protein as either a PE or PPE protein. All selected *M. spongiae* PE/PPE proteins were then aligned against known PE/PPE proteins and investigated for the presence of key domains/signature sequences according to criteria in (3).

### Putative PE proteins

| *M. spongiae* locus tag | PE motif | YXXXD/E motif | Length (AAs) | C-term Dom (larger than 111 aas) | PGRS motif (repeating GGx) | H37Rv closest match | %ID | *M. marinum* closest match | %ID | Comments | Detected by proteomics? |
|---|---|---|---|---|---|---|---|---|---|---|---|
| F6B93_00375 | Y | Y | 541 | Y | Y | RV2591 (PE_PGRS44) | 61.497 | MMAR_2113 | 68.212 | | |
| F6B93_00430 | N | Y | 1044 | Y | N | RV0101 | 37.902 | MMAR_0368 | 32.667 | PD instead of PE, YXXXD motif closer to start than other PEs | |
| F6B93_00450 | Y | Y | 549 | Y | Y | RV1452C (PE_PGRS28)c | 65.101 | MMAR_0837 | 65.079 | | |
| F6B93_00455 | Y | Y | 1489 | Y | Y | RV2591 (PE_PGRS44) | 64.667 | MMAR_1538 | 60 | | |
| F6B93_00460 | Y | Y | 1371 | Y | Y | RV1768 (PE_PGRS31) | 68.504 | MMAR_5339 | 63.492 | PE_PGRS31 involved in mycobacterial survival in macrophages (4) | Y |
| F6B93_00465 | Y | Y | 1258 | Y | Y | RV3595c (PE_PGRS59) | 65.649 | MMAR_4149 | 61.856 | | |
| F6B93_00470 | Y | Y | 1112 | Y | Y | RV1768 (PE_PGRS31) | 68.217 | MMAR_4561 | 62.205 | | |
| F6B93_00720 | Y | Y | 633 | Y | Y | RV1068c (PE_PGRS20) | 60.465 | MMAR_2103 | 61.417 | | |
| F6B93_00755 | N | Y | 2640 | Y | Y | RV3511 (PE_PGRS55) | 79.688 | MMAR_2100 | 61.983 | PD instead of PE | |
| F6B93_01010 | Y | Y | 874 | Y | Y | RV3388 (PE_PGRS52) | 56.818 | MMAR_2709 | 54.815 | | |
| F6B93_01370 | N | Y | 465 | Y | N | RV0152c (PE2) | 74.376 | MMAR_0370 | 70.172 | Truncated, possible pseudogene. PE2 is an esterase that hydrolyses p-nitrophenyl esters (5). | Y. |
| F6B93_01375 | N | Y | 593 | Y | N | RV1800 (PPE28) | 43.243 | MMAR_0369 | 55.74 | IE instead of PE | Y |
| F6B93_01420 | Y | Y | 500 | Y | N | RV0160c (PE4) | 70.06 | MMAR_0384 | 69.368 | | Y |
| F6B93_01425 | N | Y | 499 | Y | N | RV0160c (PE4) | 58.566 | MMAR_0384 | 58.185 | AD instead of PE | |
| F6B93_01480 | N | Y | 382 | Y | N | RV1184c | 50.98 | MMAR_4939 | 32.028 | SA instead of PE. Real PE?? | |
| F6B93_01780 | Y | Y | 2159 | Y | Y | RV3345c (PE_PGRS50) | 66.406 | MMAR_5339 | 66.434 | | |
| F6B93_02045 | N | Y | 607 | Y | Y | RV1768 (PE_PGRS31) | 72 | MMAR_2970 | 69.231 | PQ instead of PE | |
| F6B93_02050 | N | Y | 667 | Y | Y | RV1768 (PE_PGRS31) | 72.8 | MMAR_2933 | 56.954 | PQ instead of PE | |
| F6B93_02085 | Y | Y | 916 | Y | Y | RV0279c (PE_PGRS4) | 83.824 | MMAR_2695 | 50.714 | | |
| F6B93_02115 | Y | N | 103 | N | - | RV0285 (PE5) | 88.35 | MMAR_0544 | 82.178 | Truncated, possible pseudogene | |
| F6B93_02190 | Y | Y | 545 | Y | Y | RV3595c (PE_PGRS59) | 70.629 | MMAR_3199 | 71.918 | | |
| F6B93_02195 | Y | Y | 472 | Y | Y | RV0872c (PE_PGRS15) | 73.438 | MMAR_3199 | 69.863 | | |
| F6B93_02705 | N | N | 642 | Y | Y | RV1068c (PE_PGRS20) | 50 | MMAR_4270 | 54.545 | PD instead of PE, FXXXE instead of YXXXE/D | |

| | | | | | | | | | | | |
|---|---|---|---|---|---|---|---|---|---|---|---|
| F6B93_02960 | N | Y | 868 | Y | Y | RV0872c (PE_PGRS15) | 64.151 | MMAR_2709 | 62.992 | PQ instead of PE/PD | |
| F6B93_03025 | N | Y | 3023 | Y | Y | RV3511 (PE_PGRS55) | 79.528 | MMAR_3728 | 62.5 | PD instead of PE. | Y |
| F6B93_03225 | N | N | 36 | N | | RV0938 | 40 | MMAR_2987 | 43.333 | Probable pseudogene | |
| F6B93_03430 | Y | Y | 99 | N | | RV2396 (PE_PGRS41) | 66.327 | MMAR_2709 | 63.265 | Probable pseudogene | |
| F6B93_03885 | Y | Y | 876 | Y | Y | RV1768 (PE_PGRS31) | 63.78 | MMAR_5344 | 57.692 | | |
| F6B93_03905 | N | Y | 3935 | Y | Y | RV3595c (PE_PGRS59) | 63.559 | MMAR_1594 | 65.6 | PD instead of PE | |
| F6B93_03915 | Y | Y | 862 | Y | Y | RV1818c (PE_PGRS33) | 66.923 | MMAR_4571 | 52.381 | | |
| F6B93_04905 | N | Y | 580 | Y | Y | RV0872c (PE_PGRS15) | 66.667 | MMAR_4149 | 64.557 | AA instead of PE, has YXXXE and PGRS doms | |
| F6B93_04925 | N | Y | 99 | N | | RV3872 (PE35) | 46.939 | MMAR_0185 | 59.596 | PA instead of PE, has YXXXE motif | |
| F6B93_05215 | Y | Y | 966 | Y | Y | RV2396 (PE_PGRS41) | 83.333 | MMAR_4149 | 64.615 | | |
| F6B93_05760 | N | Y | 2486 | Y | Y | RV3511 (PE_PGRS55) | 69.6 | MMAR_4186 | 54.331 | PD instead of PE | |
| F6B93_05765 | N | Y | 1859 | Y | Y | RV1983 (PE_PGRS35) | 58.647 | MMAR_3728 | 57.971 | PD instead of PE | |
| F6B93_05770 | N | Y | 1923 | Y | Y | RV3595c (PE_PGRS59) | 60.317 | MMAR_2492 | 54.194 | PD instead of PE | |
| F6B93_05845 | Y | Y | 860 | Y | Y | RV1768 (PE_PGRS31) | 61.194 | MMAR_5344 | 57.463 | | |
| F6B93_06160 | N | Y | 669 | Y | Y | RV1452C (PE_PGRS28)c | 51.667 | MMAR_2010 | 48.406 | PG instead of PE | Y |
| F6B93_06180 | Y | Y | 409 | Y | Y | RV0872c (PE_PGRS15) | 69.531 | MMAR_4149 | 65.823 | | |
| F6B93_06300 | N | Y | 2056 | Y | Y | RV1441c (PE_PGRS26) | 73.958 | MMAR_3199 | 65.753 | PQ instead of PE/PD | |
| F6B93_06345 | Y | Y | 467 | Y | N | RV3097c (LipY) | 67.382 | MMAR_1547 | 66.667 | LipY increases Mtb virulence when overexpressed (6) | Y |
| F6B93_06355 | Y | Y | 477 | Y | Y | RV1441c (PE_PGRS26) | 66.049 | MMAR_2053 | 67.5 | | |
| F6B93_06500 | N | Y | 862 | Y | Y | RV0872c (PE_PGRS15) | 55.556 | MMAR_3105 | 54.839 | PG instead of PE | |
| F6B93_08475 | Y | Y | 772 | Y | Y | RV0872c (PE_PGRS15) | 62.759 | MMAR_1199 | 61.719 | | |
| F6B93_08690 | Y | Y | 537 | Y | Y | RV2591 (PE_PGRS44) | 60.753 | MMAR_2113 | 65.563 | | |
| F6B93_08695 | Y | Y | 540 | Y | Y | RV2591 (PE_PGRS44) | 65.591 | MMAR_2113 | 64.238 | | |
| F6B93_08715 | Y | Y | 545 | Y | Y | RV1068c (PE_PGRS20) | 60.417 | MMAR_3199 | 64.052 | | |
| F6B93_08725 | Y | Y | 1355 | Y | Y | RV3595c (PE_PGRS59) | 74.194 | MMAR_3728 | 64.384 | | |
| F6B93_08735 | Y | Y | 865 | Y | Y | RV0872c (PE_PGRS15) | 62.205 | MMAR_2100 | 60 | | |
| F6B93_08820 | N | Y | 1338 | Y | Y | RV1983 (PE_PGRS35) | 59.524 | MMAR_3199 | 69.93 | LE instead of PE | |
| F6B93_09215 | y | Y | 620 | Y | N | RV1430 (PE16) | 57.742 | MMAR_2235 | 59.734 | | Y |
| F6B93_09315 | Y | Y | 597 | Y | Y | RV1768 (PE_PGRS31) | 63.014 | MMAR_5135 | 59.155 | | |
| F6B93_09550 | N | Y | 496 | Y | Y | RV0872c (PE_PGRS15) | 65.873 | MMAR_4149 | 65.19 | PQ instead of PE | |
| F6B93_09600 | Y | Y | 742 | Y | Y | RV1430 (PE16) | 58.416 | MMAR_0383 | 39.206 | PE16 is an esterase (7). | Y |
| F6B93_09945 | Y | Y | 929 | Y | Y | RV1450c (PE_PGRS27) | 65.574 | MMAR_4186 | 66.667 | | |
| F6B93_10255 | N | Y | 372 | Y | N | RV1452c (PE_PGRS28) | 50 | MMAR_2453 | 38.108 | PD instead of PE | |
| F6B93_10260 | N | Y | 333 | Y | N | RV1646 (PE17) | 60.245 | MMAR_2453 | 52.239 | PD instead of PE | |
| F6B93_10285 | N | Y | 492 | Y | N | RV1646 (PE17) | 55.039 | MMAR_2973 | 56.034 | PD instead of PE. PE17 promotes host cell apoptosis in *M. smegmatis* (8). Enhances *M. smegmatis* survival in | Y |

| | | | | | | | | | | | |
|---|---|---|---|---|---|---|---|---|---|---|---|
| | | | | | | | | | | macrophages and pathogenicity in mice (9) | |
| F6B93_10290 | y | Y | 849 | Y | Y | RV1651c (PE_PGRS30) | 66.129 | MMAR_3570 | 49 | | |
| F6B93_10625 | Y | Y | 489 | Y | N | RV3345c (PE_PGRS50) | 58.871 | MMAR_3402 | 73.878 | | |
| F6B93_10635 | Y | Y | 859 | Y | Y | RV1768 (PE_PGRS31) | 64.567 | MMAR_4561 | 60.156 | | |
| F6B93_10760 | Y | Y | 937 | Y | Y | RV0872c (PE_PGRS15) | 66.667 | MMAR_2709 | 68.116 | | |
| F6B93_10770 | Y | Y | 921 | Y | Y | RV0872c (PE_PGRS15) | 63.492 | MMAR_2709 | 66.667 | | |
| F6B93_10840 | y | Y | 103 | N | - | RV0285 (PE5) | 84.466 | MMAR_0544 | 79.208 | Truncated, possible pseudogene | |
| F6B93_10935 | Y | Y | 628 | Y | Y | RV2396 (PE_PGRS41) | 70.079 | MMAR_4548 | 67.969 | | |
| F6B93_10975 | y | Y | 1833 | Y | Y | RV3595c (PE_PGRS59) | 62.5 | MMAR_2492 | 60.69 | | |
| F6B93_11030 | Y | Y | 100 | N | N | RV1791 (PE19) | 87 | MMAR_2670 | 90.805 | | |
| F6B93_11045 | y | Y | 100 | N | - | RV1791 (PE19) | 95 | MMAR_2673 | 97 | | |
| F6B93_11240 | y | Y | 928 | Y | Y | RV0872c (PE_PGRS15) | 67.46 | MMAR_4149 | 68.548 | | |
| F6B93_11245 | N | Y | 560 | Y | Y | RV3595c (PE_PGRS59) | 71.318 | MMAR_2709 | 73.77 | PD instead of PE | |
| F6B93_11875 | N | Y | 218 | Y | N | RV3872 (PE35) | 57.576 | MMAR_2894 | 70.732 | PS instead of PE | |
| F6B93_11935 | y | Y | 557 | Y | Y | RV0977 (PE_PGRS16) | 50.781 | MMAR_2933 | 65.436 | | Y |
| F6B93_11995 | N | Y | 468 | Y | Y | RV2853 (PE_PGRS48) | 62.59 | MMAR_2053 | 66.197 | PD instead of PE | |
| F6B93_12220 | Y | Y | 581 | Y | N | RV0152c (PE2) | 56.548 | MMAR_0370 | 61.938 | | Y |
| F6B93_12270 | N | Y | 591 | Y | Y | RV0578c (PE_PGRS7) | 62.295 | MMAR_0943 | 57.325 | | |
| F6B93_12275 | Y | Y | 858 | Y | Y | RV0872c (PE_PGRS15) | 59.524 | MMAR_2948 | 71.642 | | |
| F6B93_12285 | N | Y | 477 | Y | Y | RV2853 (PE_PGRS48) | 48.837 | MMAR_3402 | 71.992 | SD instead of PE | |
| F6B93_12365 | N | Y | 734 | Y | Y | RV3595c (PE_PGRS59) | 60.15 | MMAR_4186 | 65.385 | PQ instead of PE | |
| F6B93_12370 | N | Y | 730 | Y | Y | RV1441c (PE_PGRS26) | 68.657 | MMAR_3199 | 62.162 | PQ instead of PE | |
| F6B93_12410 | Y | Y | 914 | Y | Y | RV1651c (PE_PGRS30) | 56.627 | MMAR_2459 | 48.201 | | Y |
| F6B93_12585 | Y | Y | 524 | Y | Y | RV1983 (PE_PGRS35) | 62.308 | MMAR_3199 | 61.486 | | |
| F6B93_12665 | Y | Y | 675 | Y | Y | RV1441c (PE_PGRS26) | 68.657 | MMAR_2053 | 67.5 | | |
| F6B93_12670 | Y | Y | 623 | Y | Y | RV1441c (PE_PGRS26) | 70.349 | MMAR_2053 | 67 | | |
| F6B93_12745 | y | Y | 148 | Y | N | N/A | N/A | MMAR_4186 | 51.145 | | |
| F6B93_12750 | Y | Y | 1952 | Y | Y | N/A | N/A | MMAR_5362 | 60 | | |
| F6B93_12775 | N | N | 72 | N | N | N/A | N/A | MMAR_4818 | 46.875 | Pseudogene, C-term only of a PE or PPE prot | |
| F6B93_12785 2 | N | N | N/A | N/A | N | N/A | N/A | #N/A | #N/A | Pseudogene, C-term only of a PE or PPE prot | |
| F6B93_12790 | Y | Y | 168 | Y | N | RV1768 (PE_PGRS31) | 68.217 | MMAR_4548 | 63.359 | | |
| F6B93_12795 | Y | Y | 538 | Y | Y | RV0872c (PE_PGRS15) | 63.636 | MMAR_4149 | 60.317 | | |
| F6B93_13770 | Y | Y | 586 | Y | Y | RV3595c (PE_PGRS59) | 72.951 | MMAR_2973 | 57.983 | | |
| F6B93_13790 | Y | Y | 1985 | Y | Y | RV3511 (PE_PGRS55) | 75 | MMAR_1594 | 68.75 | | |
| F6B93_13800 | Y | Y | 5670 | Y | Y | RV2853 (PE_PGRS48) | 45.833 | MMAR_2973 | 55.085 | | |
| F6B93_13815 | Y | Y | 94 | N | N | RV2853 (PE_PGRS48) | 57.447 | MMAR_2102 | 59.574 | | |

| | | | | | | | | | | | |
|---|---|---|---|---|---|---|---|---|---|---|---|
| F6B93_13890 | y | Y | 499 | Y | Y | RV0872c (PE_PGRS15) | 69.048 | MMAR_4149 | 66.667 | | |
| F6B93_13920 | Y | Y | 1174 | Y | Y | RV0872c (PE_PGRS15) | 62.5 | MMAR_2709 | 63.333 | | |
| F6B93_14090 | N | N | 660 | Y | Y | RV0977 (PE_PGRS16) | 60.366 | MMAR_2103 | 74.316 | PD instead of PE. | Y |
| F6B93_14135 | y | Y | 839 | Y | Y | RV0278c (PE_PGRS3) | 85.106 | MMAR_4560 | 63.71 | | |
| F6B93_14145 | Y | Y | 530 | Y | Y | RV0280 (PPE3) | 63.168 | MMAR_1418 | 58.444 | | |
| F6B93_14535 | Y | Y | 866 | Y | Y | RV1818c (PE_PGRS33) | 65.185 | MMAR_4399 | 62.838 | | |
| F6B93_14545 | Y | Y | 898 | Y | Y | RV0872c (PE_PGRS15) | 66.667 | MMAR_4149 | 67.2 | | |
| F6B93_14555 | Y | Y | 867 | Y | Y | RV0872c (PE_PGRS15) | 62.162 | MMAR_4560 | 62.992 | | |
| F6B93_14590 | Y | Y | 692 | Y | Y | RV2396 (PE_PGRS41) | 62.698 | MMAR_3199 | 64.901 | | |
| F6B93_14865 | Y | N | 3625 | Y | Y | RV0355c (PPE8) | 56.044 | MMAR_0642 | 40.861 | Has YXXXL instead of YXXXE/D. | Y |
| F6B93_15090 | Y | Y | 527 | Y | Y | RV2853 (PE_PGRS48) | 64 | MMAR_4149 | 65.6 | | |
| F6B93_15150 | Y | Y | 545 | Y | Y | RV3595c (PE_PGRS59) | 66.393 | MMAR_3728 | 67.114 | | |
| F6B93_15380 | Y | Y | 1161 | Y | Y | RV1441c (PE_PGRS26) | 68.657 | MMAR_2053 | 71.831 | | |
| F6B93_15385 | Y | Y | 183 | Y | N | RV0872c (PE_PGRS15) | 63.636 | MMAR_4116 | 67.647 | | |
| F6B93_15450 | N | Y | 779 | Y | Y | RV2853 (PE_PGRS48) | 55.714 | MMAR_4149 | 62.827 | PA instead of PE | |
| F6B93_15455 | N | Y | 483 | Y | Y | RV3595c (PE_PGRS59) | 65.734 | MMAR_3199 | 64.384 | PD instead of PE | |
| F6B93_15460 | Y | N | 865 | Y | Y | RV0872c (PE_PGRS15) | 62.5 | MMAR_4399 | 64.964 | FXXXE instead of YXXXE | |
| F6B93_15465 | y | Y | 881 | Y | Y | RV1441c (PE_PGRS26) | 61.069 | MMAR_5044 | 60.769 | | |
| F6B93_15470 | Y | Y | 453 | Y | Y | RV0872c (PE_PGRS15) | 64.286 | MMAR_3199 | 64.189 | | Y |
| F6B93_15485 | Y | Y | 649 | Y | Y | RV3595c (PE_PGRS59) | 61.972 | MMAR_2053 | 68.098 | | |
| F6B93_15630 | N | N | 28 | N | | RV1768 (PE_PGRS31) | 69.565 | MMAR_2053 | 78.571 | Pseudogene, C-term only of a PE or PPE prot | |
| F6B93_15635 | Y | Y | 593 | Y | Y | RV1441c (PE_PGRS26) | 69.347 | MMAR_4186 | 67.969 | | |
| F6B93_15705 | y | Y | 651 | Y | Y | RV1452C (PE_PGRS28)c | 63.636 | MMAR_4548 | 66.412 | | Y |
| F6B93_15710 | N | Y | 95 | N | N | RV3097c (LipY) | 48.421 | MMAR_2973 | 53.684 | PD instead of PE | |
| F6B93_15720 | N | Y | 886 | Y | Y | RV1803c (PE_PGRS32) | 78.261 | MMAR_5339 | 53.543 | PV instead of PE | |
| F6B93_15815 | Y | Y | 794 | Y | Y | RV1441c (PE_PGRS26) | 61.364 | MMAR_4149 | 60.156 | | |
| F6B93_15920 | y | Y | 619 | Y | Y | RV1768 (PE_PGRS31) | 68.548 | MMAR_5135 | 60.284 | | |
| F6B93_16160 | y | Y | 2565 | Y | Y | RV2162c (PE_PGRS38) | 70.732 | MMAR_0943 | 57.851 | | |
| F6B93_16175 | N | N | 1449 | Y | Y | RV1864c | 46.667 | N/A | N/A | Start after PE dom, GXXXG instead of YXXXE, but has PGRS dom | |
| F6B93_16280 | Y | Y | 444 | Y | Y | RV0872c (PE_PGRS15) | 65.625 | MMAR_4149 | 64.78 | | |
| F6B93_16285 | Y | Y | 682 | Y | Y | RV1325c (PE_PGRS24) | 68.548 | MMAR_4149 | 67.089 | | |
| F6B93_16500 | Y | Y | 647 | Y | Y | RV1441c (PE_PGRS26) | 69.291 | MMAR_5339 | 65.354 | | |
| F6B93_16505 | Y | Y | 686 | Y | Y | RV2853 (PE_PGRS48) | 52.308 | MMAR_2010 | 54.225 | | |
| F6B93_17035 | y | Y | 584 | Y | Y | RV2853 (PE_PGRS48) | 64.171 | MMAR_1208 | 60.163 | | |
| F6B93_17090 | N | Y | 1520 | Y | Y | RV0977 (PE_PGRS16) | 47.748 | MMAR_4278 | 57.788 | AQ instead of PE | |
| F6B93_17110 | y | Y | 289 | Y | N | RV1172c (PE12) | 56.61 | MMAR_4280 | 61.071 | | |

| | | | | | | | | | | | |
|---|---|---|---|---|---|---|---|---|---|---|---|
| F6B93_17350 | N | Y | 605 | Y | N | RV1800 (PPE28) | 49.677 | MMAR_3984 | 53.984 | PQ instead of PE. PPE28 helps to evade intracellular killing in *M. smegmatis* (10) | Y |
| F6B93_17510 | N | Y | 924 | Y | Y | RV0872c (PE_PGRS15) | 60.769 | MMAR_5339 | 61.417 | PD instead of PE | |
| F6B93_17530 | N | N | 97 | N | | RV1452C (PE_PGRS28) | 56.842 | MMAR_4186 | 56.701 | PA instead of PE and LxxxE instead of YxxxE. Pseudogene? | |
| F6B93_17645 | N | Y | 616 | Y | N | RV0754 (PE_PGRS11) | 66.34 | MMAR_4953 | 62.541 | PD instead of PE | Y |
| F6B93_17690 | N | Y | 875 | Y | Y | RV0746 (PE_PGRS9) | 82.812 | MMAR_2948 | 65 | PQ instead of PE | |
| F6B93_17700 | y | Y | 273 | Y | N | RV1040c (PE8) | 74.101 | MMAR_4451 | 74.368 | | |
| F6B93_17960 | N | Y | 675 | Y | Y | RV0872c (PE_PGRS15) | 51.493 | MMAR_2709 | 51.111 | PQ instead of PE | |
| F6B93_18055 | Y | Y | 1160 | Y | Y | RV2487c (PE_PGRS42) | 56.41 | MMAR_2656 | 55.634 | | |
| F6B93_18060 | Y | Y | 1237 | Y | N | RV1452C (PE_PGRS28)c | 54.483 | MMAR_3199 | 49.664 | | |
| F6B93_18090 | y | Y | 1256 | Y | Y | RV1441c (PE_PGRS26) | 62.353 | MMAR_2053 | 60.494 | | Y |
| F6B93_18095 | N | Y | 564 | Y | N | RV0152c (PE2) | 67.322 | MMAR_3427 | 73.451 | PQ instead of PE | Y |
| F6B93_18100 | N | N | 99 | N | N | RV0872c (PE_PGRS15) | 62.857 | MMAR_2053 | 68.571 | Pseudogene | |
| F6B93_18195 | Y | N | 255 | Y | N | RV2853 (PE_PGRS48) | 59.2 | MMAR_2695 | 57.692 | YXXXK instead of YXXXE | |
| F6B93_18205 | Y | Y | 188 | Y | N | RV0872c (PE_PGRS15) | 66.406 | MMAR_3199 | 64.189 | | |
| F6B93_18395 | y | Y | 322 | Y | Y | RV0872c (PE_PGRS15) | 54.962 | MMAR_4548 | 58.779 | | |
| F6B93_18400 | Y | Y | 577 | Y | Y | RV0872c (PE_PGRS15) | 61.111 | MMAR_2709 | 65.041 | | |
| F6B93_18430 | Y | Y | 1270 | Y | Y | RV2487c (PE_PGRS42) | 64.179 | MMAR_2656 | 55.072 | | |
| F6B93_18480 | N | Y | 534 | Y | Y | RV0872c (PE_PGRS15) | 63.194 | MMAR_4149 | 62.025 | PD instead of PE | |
| F6B93_18485 | Y | Y | 157 | Y | N | RV0872c (PE_PGRS15) | 62.281 | MMAR_3199 | 60.526 | | |
| F6B93_18635 | N | Y | 897 | Y | Y | RV2853 (PE_PGRS48) | 75.862 | MMAR_2645 | 70.909 | TD instead of PE | |
| F6B93_18645 | Y | N | 51 | N | N | RV0916c (PE7) | 57.143 | MMAR_4995 | 52 | Pseudogene | |
| F6B93_19045 | Y | Y | 1351 | Y | Y | RV1452C (PE_PGRS28)c | 62.5 | MMAR_3199 | 65.359 | | |
| F6B93_19065 | Y | Y | 845 | Y | Y | RV1452C (PE_PGRS28)c | 65.278 | MMAR_3199 | 64.238 | | |
| F6B93_19170 | Y | Y | 2175 | Y | Y | RV3511 (PE_PGRS55) | 71.094 | MMAR_4270 | 58.451 | | |
| F6B93_19185 | N | N | 1872 | Y | Y | N/A | N/A | N/A | N/A | Pseudogene | |
| F6B93_19310 | Y | Y | 125 | Y | N | RV3097c (LipY) | 43.925 | MMAR_0625 | 54.762 | | |
| F6B93_19435 | N | Y | 1073 | Y | Y | RV2853 (PE_PGRS48) | 55.385 | MMAR_2709 | 50.704 | PA instead of PE | |
| F6B93_19440 | N | Y | 1455 | Y | Y | RV2853 (PE_PGRS48) | 57.513 | MMAR_2709 | 52.083 | PG instead of PE | |
| F6B93_19445 | N | Y | 1274 | Y | Y | RV2490c (PE_PGRS43) | 54.487 | MMAR_3199 | 50.676 | PA instead of PE | |
| F6B93_19450 | N | Y | 211 | Y | Y | RV2853 (PE_PGRS48) | 56.477 | MMAR_2709 | 52.083 | PG instead of PE | |
| F6B93_19455 | N | Y | 1912 | Y | Y | RV1768 (PE_PGRS31) | 51.163 | MMAR_4560 | 58.14 | PA instead of PE | |
| F6B93_19805 | Y | Y | 502 | Y | Y | RV3595c (PE_PGRS59) | 67.669 | MMAR_3199 | 69.595 | | |
| F6B93_19830 | Y | Y | 1939 | Y | Y | RV0977 (PE_PGRS16) | 59.722 | MMAR_2248 | 66 | | |
| F6B93_19870 | Y | Y | 852 | Y | Y | RV1768 (PE_PGRS31) | 66.142 | MMAR_2695 | 65.278 | | |
| F6B93_20180 | N | Y | 1428 | Y | Y | RV0832 (PE_PGRS12) | 57.364 | MMAR_4149 | 50.92 | PQ instead of PE | |
| F6B93_20395 | Y | Y | 1455 | Y | Y | RV0872c (PE_PGRS15) | 61.111 | MMAR_2709 | 63.78 | *M. spongiae* orthologue looks | |

| | | | | | | | | | | | |
|---|---|---|---|---|---|---|---|---|---|---|---|
| | | | | | | | | | | to be intact cf Mtb | |
| F6B93_20475 | n | Y | 537 | Y | Y | RV3388 (PE_PGRS52) | 65.854 | MMAR_3728 | 62.838 | PQ instead of PE/PD. PE_PGRS52 orthologue | Y |
| F6B93_20815 | Y | Y | 100 | N | N | RV3622c (PE32) | 75.281 | MMAR_5122 | 71.91 | | |
| F6B93_20825 | y | y | 100 | N | - | RV3622c (PE32) | 75.862 | MMAR_5122 | 72.414 | | |
| F6B93_20935 | Y | Y | 856 | Y | Y | RV1818c (PE_PGRS33) | 73.016 | MMAR_2709 | 63.78 | | |
| F6B93_21290 | y | Y | 1421 | Y | N | RV3595c (PE_PGRS59) | 67.46 | MMAR_3728 | 60.959 | | |
| F6B93_21460 | N | Y | 571 | Y | Y | RV1768 (PE_PGRS31) | 61.417 | MMAR_4399 | 70.213 | PD instead of PE | |
| F6B93_21525 | N | Y | 564 | Y | N | RV0152c (PE2) | 67.076 | MMAR_3427 | 73.451 | PQ instead of PE | |
| F6B93_21725 | N | Y | 795 | Y | Y | RV1441c (PE_PGRS26) | 69.136 | MMAR_2053 | 69.753 | PQ instead of PE | |
| F6B93_21985 | y | Y | 571 | Y | N | RV3812 (PE_PGRS62) | 58.111 | MMAR_2460 | 43.213 | | |
| F6B93_21990 | Y | Y | 562 | Y | N | RV3812 (PE_PGRS62) | 63.377 | MMAR_2460 | 39.089 | PE_PGRS62 promotes survival of *M. smegmatis* in macrophages (11). Blocks phagosome maturation in *M. smegmatis* and *M. marinum* (12). Antibodies to PE_PGRS62 associated with latent TB (13). | Y |
| F6B93_22305 | N | N | 172 | Y | N | RV3372A | 69.565 | MMAR_0299 | 62.069 | Pseudogene, C-term only of a PE or PPE prot | |
| F6B93_22310 | N | Y | 284 | Y | N | RV0872c (PE_PGRS15) | 58.163 | MMAR_2709 | 64.286 | AE instead of PE and YXXXD instead of YXXXE | |
| F6B93_22315 | n | Y | 552 | Y | N | RV0152c (PE2) | 63.99 | MMAR_0370 | 63.225 | PQ instead of PE/PD | |
| F6B93_22390 | y | Y | 101 | N | - | RV3893c (PE36) | 86.842 | MMAR_1949 | 34.343 | | |

### Putative PPE proteins

| *M. spongiae* locus tag | PPE motif | WxG motif | Amino acids | C-term Dom (larger than200aas) | PxxPxxW motif (10-30 aas from C-term) | SVP motif (near C-term) | NxGxGNxG repeats | H37Rv orthologue | %ID | *M. marinum* orthologue | %ID | Comments | Detected by proteomics? |
|---|---|---|---|---|---|---|---|---|---|---|---|---|---|
| F6B93_00200 | Y | Y | 400 | Y | N | Y | N | RV1706c (PPE23) | 53.431 | MMAR_2517 | 46.348 | | |
| F6B93_00205 | Y | Y | 390 | Y | N | Y | N | RV1705c (PPE22) | 64.031 | MMAR_2516 | 55.051 | | |
| F6B93_01055 | Y | Y | 508 | Y | Y | N | N | RV0286 (PPE4) | 52.574 | MMAR_1418 | 40.952 | | |
| F6B93_01065 | Y | Y | 483 | Y | Y | N | N | RV0096 (PPE1) | 62.422 | MMAR_0261 | 55.67 | | |
| F6B93_02090 | Y | Y | 559 | Y | Y | N | N | RV0280 (PPE3) | 64.545 | MMAR_1418 | 57.168 | | |
| F6B93_02120 | Y | Y | 524 | Y | Y | N | N | RV0286 (PPE4) | 67.113 | MMAR_0545 | 63.527 | | |
| F6B93_02510 | Y | Y | 3602 | Y | N | N | Y | RV0355c (PPE8) | 77.368 | MMAR_0642 | 66.225 | | Y |
| F6B93_02720 | Y | Y | 438 | Y | N | N | N | RV0387c | 68.1 | MMAR_0685 | 64.189 | | |
| F6B93_02750 | Y | Y | 414 | Y | N | Y | N | RV1361c (PPE19) | 40.625 | MMAR_4240 | 42.217 | | |

| | | | | | | | | | | | | | |
|---|---|---|---|---|---|---|---|---|---|---|---|---|---|
| F6B93_03045 | Y | Y | 500 | Y | N | N | Y | RV0442c (PPE10) | 70.226 | MMAR_0761 | 65.039 | | |
| F6B93_03390 | Y | Y | 406 | Y | N | Y | N | RV3532 (PPE61) | 55.288 | MMAR_3444 | 49.148 | | |
| F6B93_03445 | Y | Y | 2276 | Y | N | Y | Y | RV1918c (PPE35) | 46.527 | MMAR_1028 | 47.17 | | |
| F6B93_03515 | Y | Y | 536 | Y | N | N | Y | RV0355c (PPE8) | 58.75 | MMAR_0642 | 56.569 | | Y |
| F6B93_04640 | Y | Y | 366 | Y | N | Y | N | RV1705c (PPE22) | 57.254 | MMAR_1094 | 68.241 | | |
| F6B93_04645 | Y | Y | 366 | Y | N | N | N | RV1790 (PPE27) | 44.321 | MMAR_1095 | 55.026 | | |
| F6B93_04930 | Y | Y | 370 | Y | N | N | N | RV3873 (PPE68) | 59.444 | MMAR_0186 | 61.157 | | |
| F6B93_05085 | Y | Y | 3739 | Y | N | Y | Y | RV3350c (PPE56) | 68.116 | MMAR_0642 | 39.974 | | |
| F6B93_05155 | N | Y | 465 | Y | N | N | N | RV3539 (PPE63) | 58.7 | MMAR_4939 | 54.882 | RPE instead of PPE. Promotes macrophage survivale when expressed in *M. smegmatis* (14). | Y |
| F6B93_05255 | Y | Y | 864 | Y | N | N | Y | RV0355c (PPE8) | 51.429 | MMAR_3546 | 52.595 | | |
| F6B93_05300 | Y | Y | 380 | Y | N | N | N | RV1705c (PPE22) | 40.816 | MMAR_1235 | 67.895 | | |
| F6B93_05850 | Y | Y | 923 | Y | N | Y | Y | RV1753c (PPE24) | 51.74 | MMAR_2822 | 51.066 | NxGxGNxG motif is more towards middle of protein. | |
| F6B93_06005 | Y | Y | 621 | Y | N | N | Y | RV3159c (PPE53) | 64.687 | MMAR_1402 | 68.132 | | |
| F6B93_06010 | Y | Y | 1130 | Y | N | N | Y | RV3533c (PPE62) | 61.175 | MMAR_4320 | 55.207 | | |
| F6B93_06090 | Y | Y | 435 | Y | N | N | N | RV3144c (PPE52) | 88.068 | MMAR_1484 | 85.227 | | |
| F6B93_06205 | Y | Y | 381 | Y | N | Y | N | RV3136 (PPE51) | 66.329 | MMAR_0191 | 64.781 | | |
| F6B93_06210 | Y | Y | 371 | Y | N | Y | N | RV3136 (PPE51) | 66.237 | MMAR_0191 | 64.136 | | |
| F6B93_06215 | Y | Y | 378 | Y | N | Y | N | RV3136 (PPE51) | 62.564 | MMAR_0191 | 61.88 | | |
| F6B93_06605 | Y | Y | 1025 | Y | N | Y | Y | RV1753c (PPE24) | 56.932 | MMAR_2822 | 57.019 | PPE24 orthologue | Y |
| F6B93_07240 | Y | Y | 364 | Y | N | Y | N | RV2352c (PPE38) | 47.041 | MMAR_4324 | 48.974 | | |
| F6B93_07490 | Y | Y | 399 | Y | N | Y | N | RV1808 (PPE32) | 48.246 | MMAR_1847 | 56.436 | | |
| F6B93_07775 | Y | Y | 986 | Y | N | Y | Y | RV1753c (PPE24) | 54.394 | MMAR_2822 | 53.853 | | |
| F6B93_07855 | Y | Y | 381 | Y | N | N | N | RV2770c (PPE44) | 65.455 | MMAR_1947 | 61.68 | | |
| F6B93_08245 | Y | Y | 1227 | Y | N | N | Y | RV3533c (PPE62) | 43.415 | MMAR_3546 | 54.698 | | |
| F6B93_08595 | Y | Y | 603 | Y | N | N | Y | RV2608 (PPE42) | 72.297 | MMAR_2235 | 50.987 | | |
| F6B93_09260 | Y | Y | 1121 | Y | N | N | Y | RV1135c (PPE16) | 61.396 | MMAR_4320 | 57.725 | | |
| F6B93_09265 | Y | Y | 1133 | Y | N | N | Y | RV2356c (PPE40) | 65.909 | MMAR_4320 | 57.407 | | Y |
| F6B93_09270 | Y | Y | 1102 | Y | N | N | Y | RV1135c (PPE16) | 61.605 | MMAR_4320 | 59.067 | | |
| F6B93_09830 | Y | N | 119 | N | N | N | N | RV1800 (PPE28) | 84.211 | MMAR_2822 | 80 | VxG instead of WxG, possibly truncated | |
| F6B93_10835 | Y | Y | 602 | Y | Y | N | N | RV0256c (PPE2) | 70.859 | MMAR_2944 | 61.888 | | |
| F6B93_10905 | Y | Y | 612 | Y | N | Y | N | RV1809 (PPE33) | 46.361 | MMAR_0191 | 40.704 | | |
| F6B93_11025 | Y | Y | 419 | Y | N | Y | N | RV1787 (PPE25) | 49.762 | MMAR_2669 | 75.472 | | |

| | | | | | | | | | | | | | |
|---|---|---|---|---|---|---|---|---|---|---|---|---|---|
| F6B93_11035 | Y | Y | 406 | Y | N | N | N | RV1789 (PPE26) | 52.709 | MMAR_2671 | 72.263 | | |
| F6B93_11085 | Y | Y | 382 | Y | N | Y | N | RV1807 (PPE31) | 67.157 | MMAR_2683 | 61.22 | | |
| F6B93_11090 | Y | Y | 429 | Y | N | Y | N | RV1808 (PPE32) | 59.294 | MMAR_1461 | 65.046 | | |
| F6B93_11095 | Y | Y | 469 | Y | N | Y | N | RV1802 (PPE30) | 50.74 | MMAR_2685 | 69.149 | PPE30 induces macrophage apoptosis/cell death (15), suppresses proinflammatory immune response (16) | Y |
| F6B93_11660 | Y | Y | 977 | Y | N | Y | Y | RV1918c (PPE35) | 66.107 | MMAR_2822 | 60.439 | | |
| F6B93_11735 | Y | Y | 1473 | Y | N | N | Y | RV3558 (PPE64) | 62.995 | MMAR_1639 | 41.324 | | |
| F6B93_12230 | Y | Y | 3494 | Y | N | N | Y | RV0355c (PPE8) | 67.701 | MMAR_0642 | 65.181 | | |
| F6B93_12265 | N | Y | 3159 | Y | Y | Y | Y | RV0304c (PPE5) | 66.175 | MMAR_1639 | 53.658 | APE instead of PPE | |
| F6B93_12655 | Y | Y | 925 | Y | N | N | Y | N/A | N/A | MMAR_2822 | 48.677 | | |
| F6B93_12675 | Y | Y | 1135 | Y | N | N | Y | RV1135c (PPE16) | 65.075 | MMAR_4320 | 62.306 | | |
| F6B93_12765 | Y | Y | 1192 | Y | N | N | Y | N/A | N/A | MMAR_1028 | 50.282 | | |
| F6B93_13000 | Y | Y | 970 | Y | N | N | Y | RV1753c (PPE24) | 52.548 | MMAR_2822 | 50.574 | | |
| F6B93_13360 | Y | Y | 454 | Y | N | N | N | RV1809 (PPE33) | 45.385 | MMAR_2928 | 46.868 | | |
| F6B93_13365 | Y | Y | 470 | Y | N | Y | N | RV1801 (PPE29) | 44.903 | MMAR_2928 | 50.316 | | |
| F6B93_13380 | Y | Y | 513 | Y | Y | N | N | RV0286 (PPE4) | 58.175 | MMAR_1418 | 46.225 | | |
| F6B93_14130 | Y | N | 49 | N | | | | RV0256c (PPE2) | 91.667 | MMAR_2944 | 72.414 | pseudogene | |
| F6B93_14560 | Y | Y | 1888 | Y | N | N | Y | RV3558 (PPE64) | 62.995 | MMAR_1639 | 40.114 | | |
| F6B93_14895 | Y | Y | 395 | Y | N | Y | N | RV2352c (PPE38) | 70.864 | MMAR_3661 | 68.657 | | |
| F6B93_14900 | Y | Y | 398 | Y | N | Y | N | RV2352c (PPE38) | 67.654 | MMAR_3661 | 65.926 | | |
| F6B93_14905 | Y | Y | 640 | Y | N | N | Y | RV2356c (PPE40) | 65.102 | MMAR_3665 | 72.118 | | |
| F6B93_14910 | Y | Y | 626 | Y | N | N | Y | RV2356c (PPE40) | 74.722 | MMAR_3666 | 69.984 | | |
| F6B93_15785 | Y | Y | 1078 | Y | N | N | Y | RV1135c (PPE16) | 71.803 | MMAR_4320 | 62.764 | | |
| F6B93_15820 | Y | Y | 766 | Y | N | Y | Y | RV1753c (PPE24) | 56.765 | MMAR_1028 | 52.452 | | |
| F6B93_16155 | Y | N | 159 | N | N | N | N | RV0355c (PPE8) | 59.459 | MMAR_1129 | 66.234 | ExG instead of WxG | |
| F6B93_16895 | Y | Y | 1334 | Y | N | N | Y | RV1135c (PPE16) | 54.234 | MMAR_4320 | 51.192 | | |
| F6B93_17260 | Y | Y | 1182 | Y | N | N | Y | RV1135c (PPE16) | 62.946 | MMAR_4320 | 61.648 | | |
| F6B93_17265 | Y | Y | 1144 | Y | N | N | Y | RV2356c (PPE40) | 71.023 | MMAR_4320 | 60 | | |
| F6B93_17275 | Y | Y | 395 | Y | N | Y | N | RV2352c (PPE38) | 70.37 | MMAR_3661 | 68.408 | | |
| F6B93_17705 | Y | Y | 397 | Y | N | N | N | RV1039c (PPE15) | 72.613 | MMAR_4452 | 70.781 | | |
| F6B93_18140 | Y | Y | 90 | N | N | N | N | RV3159c (PPE53) | 72.881 | MMAR_4554 | 69.492 | Probably pseudogene | |
| F6B93_18650 | Y | Y | 417 | Y | N | Y | N | RV0915c (PPE14) | 64.706 | MMAR_2534 | 59.569 | | |
| F6B93_18685 | Y | Y | 3001 | Y | N | N | Y | RV0355c (PPE8) | 66.184 | MMAR_4621 | 69.127 | | |
| F6B93_19040 | N | Y | 3753 | Y | N | Y | Y | RV3350c (PPE56) | 65.928 | MMAR_0642 | 35.955 | APE instead of PPE | |
| F6B93_19055 | Y | Y | 3754 | Y | N | Y | N | RV3350c (PPE56) | 65.918 | MMAR_0642 | 38.66 | | |

| | | | | | | | | | | | | |
|---|---|---|---|---|---|---|---|---|---|---|---|---|
| F6B93_19120 | N | Y | 600 | Y | N | N | Y | RV3159c (PPE53) | 62.189 | MMAR_1469 | 57.006 | |
| F6B93_20255 | Y | Y | 754 | Y | N | N | Y | RV1135c (PPE16) | 50.307 | MMAR_4218 | 70 | |
| F6B93_20810 | Y | Y | 416 | Y | N | Y | N | RV3621c (PPE65) | 74.101 | MMAR_5121 | 67.703 | |
| F6B93_20820 | Y | Y | 417 | Y | N | Y | N | RV3621c (PPE65) | 73.923 | MMAR_5121 | 68.019 | |
| F6B93_21295 | Y | Y | 1021 | Y | N | Y | Y | RV1753c (PPE24) | 51.955 | MMAR_0932 | 44.526 | |
| F6B93_22240 | Y | Y | 371 | Y | N | N | N | RV3873 (PPE68) | 73.109 | MMAR_5448 | 71.429 | |
| F6B93_22385 | N | Y | 402 | Y | N | N | N | RV3892c (PPE69) | 81.25 | MMAR_4606 | 33.333 | TPE instead of PPE |

**Table S8**. Orthologues of mycolic acid biosynthesis proteins in *M. spongiae.*

| ***M. spongiae* locus tag** | **Protein** | **Function** | **Closest match (locus tag)[a]** | **% identity** | **Identified by proteomics** |
|---|---|---|---|---|---|
| F6B93_15935 | FasI | Type I fatty acid synthase | Rv2524c | 91 | Y |
| F6B93_13955 | FabD | Malonyl-CoA:AcpM transacylase | Rv2243 | 86 | |
| F6B93_13960 | AcpM | Acyl carrier protein | Rv2244 | 92 | Y |
| F6B93_13965 | KasA | β-Ketoacyl-AcpM synthase I | Rv2245 | 93 | Y |
| F6B93_13970 | KasB | β-Ketoacyl-AcpM synthase II | Rv2246 | 92 | Y |
| F6B93_09510 | MabA | β-Ketoacyl-AcpM reductase | Rv1483 | 92 | Y |
| F6B93_09515 | InhA | 2-trans-Enoyl-AcpM reductase | Rv1484 | 95 | Y |
| F6B93_03995 | HadA | β-Hydroxyacyl-AcpM dehydratase | Rv0635 | 86 | Y |
| F6B93_04000 | HadB | β-Hydroxyacyl-AcpM dehydratase | Rv0636 | 92 | Y |
| F6B93_04005 | HadC | β-Hydroxyacyl-AcpM dehydratase | MMAR_0970 | 88 | Y |
| F6B93_03570 | FabH | β-Ketoacyl-AcpM synthase III | MMAR_0879 | 83 | |
| F6B93_13975 | AccD6 | Carboxyltransferase subunit of acetyl-CoA carboxylase | Rv2247 | 97 | Y |
| F6B93_05380 | AccD5 | Carboxyltransferase subunit of propionyl-CoA carboxylase | Rv3280 | 94 | Y |
| F6B93_05355 | AccA3 | Biotin carboxylase subunit of acetyl-, propionyl, and fatty acyl-CoA carboxylase | Rv3285 | 92 | Y |
| F6B93_04045 | MmaA1 | Methyl branch in trans-cyclopropanated, oxygenated mycolates | Rv0645c | 88 | Y |
| F6B93_10745 | MmaA2 | cis-Cyclopropane synthase of α and oxygenated mycolates | Rv0644c | 86 | Y |
| F6B93_04040 | MmaA3 | Methoxy mycolic acid synthase | MKAN_19390 | 83 | |
| F6B93_04035 | MmaA4 | Hydroxy mycolic acid synthase | Rv0642c | 91 | Y |
| F6B93_03375 | CmaA2 | trans-Cyclopropane synthase of oxygenated mycolates, cis-Cyclopropane synthase of oxygenated mycolates (redundant with mmaA2). Distal cyclopropane synthase of α-mycolates | MMAR_0831 | | Y |
| F6B93_03195 | PcaA | cis-Cyclopropane synthase of α-mycolates | Rv0470c | 89 | Y |
| F6B93_03190 | UmaA1 | Methyl transferase of unknown specificity | Rv0469 | 91 | Y |
| F6B93_21920 | AccD4 | Carboxyltransferase subunit of fatty acyl-CoA carboxylase | Rv3799c | 95 | Y |
| F6B93_21925 | Pks13 | Polyketide synthase | Rv3800c | 86 | Y |
| F6B93_21930 | FadD32 | Fatty acyl-AMP ligase/fatty acyl-ACP synthetase | Rv3801c | 92 | Y |
| F6B93_15810 | CmrA | β-Ketoacyl reductase | MMAR_3856 | 89 | Y |

[a] Rv refers to locus tags in the *M. tuberculosis* H37Rv genome (NC_018143), MMAR locus tags are from the *M. marinum M* genome (CP000854), and Mkan locus tags are from the *M. kansasii* ATCC 12478 genome (NC_022663).

**Table S9.** Summary of key lipid species identified in *M. spongiae*.

| | | No. of species |
|---|---|---|
| **No.** | **Lipid (Sub)Class** | ***M. spongiae*** |
| 1 | Phosphatidylinositol | 12 |
| 2 | Monoacylglycerophosphoinositoldimannoside (lyso-PIM2) | 0 |
| 3 | Diacylglycerophosphoinositolmonomannoside (PIM1) | 5 |
| 4 | Diacylglycerophosphoinositoldimannoside (PIM2) | 2 |
| 5 | Monoacylated-diacylglycerophosphoinositolmonomannoside (AcPIM1) | 5 |
| 6 | Monoacylated-diacylglycerophosphoinositoldimannoside (AcPIM2) | 6 |
| 7 | Monoacylated-diacylglycerophosphoinositoltrimannoside (Ac1PIM3) | 1 |
| 8 | Monoacylated-diacylglycerophosphoinositolhexamannoside (Ac1PIM6) | 1 |
| 9 | Diacylated-diacylglycerophosphoinositolmonomannoside (Ac2PIM1) | 3 |
| 10 | Diacylated-diacylglycerophosphoinositolmonomannoside (Ac2PIM2) | 3 |
| 11 | Phosphatidic acid (PA) | 12 |
| 12 | Monoacylglycerolphosphoethanolamine (lyso-PE) | 4 |
| 13 | Diacylglycerolphosphoethanolamine (PE) | 15 |
| 14 | Phosphatidylglycerol (PG) | 2 |
| 15 | Cardiolipin (CL) | 18 |
| 16 | Monoacylglycerol (MAG) | 6 |
| 17 | Diacylglycerol (DAG) | 0 |
| 18 | Triacylglycerol (TAG) | 75 |
| 19 | Cytidine diphosphate diacylglycerol (CDP-DAG) | 1 |
| 20 | Diglycosylated Diacylglycerol (DGDG) | 1 |
| 21 | α-Mycolic acid (α-MA) and Mycolic acid (undefined) | 15 |
| 22 | Trehalose monomycolate (TMM) | 8 |
| 23 | Diacyltrehalose (DAT) | 0 |
| 24 | Diacylhexose | 0 |
| 25 | Hydroxyphthioceranic Acid | 1 |
| 26 | Phthioceranic Acid | 2 |
| 27 | Monoglycosylated phenolic glycolipid (MonoPGLA) precursor | 0 |
| 28 | Monoglycosylated phenolic glycolipid (MonoPGLA / Mono(OH)PGL) | 0 |
| 29 | Monoglycosylated phenolic glycolipid (MonoPGLB) | 0 |
| 30 | Phthiocerol dimycocerosate (PDIMA or PDIM OH) | 35 |
| 31 | Phthiocerol dimycocerosate (PDIMB) | 16 |
| 32 | Mycosanoic Acid | 4 |
| 33 | Mycocerosic Acid | 1 |
| 34 | Mycobactin (MB), Monodeoxy-Mycobactin (MDM), Dideoxy-Mycobactin (DDM) | 11 |
| 35 | Phosphomykoketides | 1 |
| 36 | Menaquinone (MK; Vitamin K2) | 4 |
| 37 | Carotenes | 1 |
| | **SUM:** | 227 |

**Table S10.** Orthologues involved in the biosynthesis of phthiocerol dimycocerosates, phenolic glycolipids, and p-hydroxybenzoic acids.

| **H37Rv locus tag** | **Protein** | **Function** | ***M. spongiae* locus tag** | **% identity** | **Identified by proteomics** |
|---|---|---|---|---|---|
| Rv2928 | TesA | Type II thioesterase thought to be involved in the release of phthiocerol and phenolphthiocerol from PpsE in the biosynthesis of PDIM and PGL | F6B93_07175 | 68 | |

| | | | | | |
|---|---|---|---|---|---|
| Rv2930 | FadD26 | Long-chain fatty acyl-AMP ligase providing PpsA with activated long-chain fatty acid starter units for the generation of phthiocerol in the biosynthesis of PDIM | F6B93_07170 | 78 | |
| Rv2931 | PpsA | Type 1 polyketide synthase responsible with PpsB-C-D-E for the elongation of C22 to C24 fatty acids and p-hydroxyphenylalkanoates with malonyl-CoA and methylmalonyl-CoA to yield phthiocerol and phenolphthiocerol derivatives,respectively. | F6B93_07165 | 55 | |
| Rv2932 | PpsB | | F6B93_07160 | 48 | |
| Rv2933 | PpsC | | F6B93_07155 | 72 | Y |
| Rv2934 | PpsD | | F6B93_07150 | 69 | |
| Rv2935 | PpsE | | F6B93_07145 | 75 | Y |
| Rv2936 | DrrA | Component of the DrrABC ABC-transporter involved in the translocation of PDIM (and PGL?) across the plasma membrane; ATP-binding protein | F6B93_07175 | 78 | |
| Rv2937 | DrrB | | F6B93_07135 | 59 | |
| Rv2938 | DrrC | | F6B93_07130 | 63 | |
| Rv2939 | PapA5 | Acyltransferase responsible for the transfer of mycocerosic acids to phthiocerol to form PDIM | F6B93_07125 | 60 | |
| Rv2940c | Mas | Mycocerosic acid synthase (type I polyketide synthase) involved in the biosynthesis of PDIM and PGL | F6B93_07120 | 75 | Y |
| Rv2941 | FadD28 | Long-chain fatty acyl-AMP ligase responsible for providing the long-chain fatty acid starter unit to Mas for the generation of mycocerosic acids | F6B93_07115 | 71 | |
| Rv2942 | MmpL7 | RND superfamily inner membrane transporter involved in PDIM translocation to the periplasm (and PGL?) | F6B93_07110 | 56 | |
| Rv2945 | LppX | RND superfamily inner membrane transporter involved in PDIM translocation to the periplasm (and PGL?) | F6B93_07105 | 67 | Y |
| Rv2946c | Pks1 | Together with Pks15, type I polyketide synthase involved in the elongation of p-hydroxybenzoic acid derivatives with malonyl-CoA to form p-hydroxyphenylalkanoates (precursors of PGL) | - | - | |
| Rv2947c | Pks15 | | - | - | |
| Rv2948c | FadD22 | p-Hydroxybenzoyl-AMP ligase involved in the biosynthesis of PGL; catalyzes the activation of p-hydroxybenzoic acid and its subsequent transfer onto Pks15/1 for the production of p-hydroxyphenylalkanoates | - | - | |
| Rv2949c | | p-Hydroxybenzoic acid synthase | - | - | |
| Rv2950c | FadD29 | Fatty acyl-AMP ligase involved in the biosynthesis of PGL; catalyzes the activation of hydroxyphenylalkanoates that are then transferred onto PpsA to yield phenolphthiocerol | - | - | |
| Rv2951c | | Ketoreductase catalyzing the reduction of (phenol)phthiodiolone to yield (phenol)phthiotriol in the biosynthesis of PDIM and PGL | - | - | |
| Rv2952 | | SAM-dependent O-methyltransferase involved in the formation of (phenol) phthiocerol dimycocerosates from (phenol)phthiodiolone dimycocerosates. Catalyzes the transfer of a methyl group to the third hydroxyl group of (phenol) phthiotriol in PDIM and glycosylated phenolphthiotriol dimycocerosates | - | - | |
| Rv2953 | | Enoyl-reductase acting in concert with PpsD in the biosynthesis of the (phenol)phthiocerol moiety of PDIM and PGL | F6B93_07095 | 70 | |
| Rv2954c | | SAM-dependent methyltransferase responsible for the O-methylation of the hydroxyl group at position 3 of the fucosyl residue of PGL (and possibly p-HBAD) | - | - | |
| Rv2955c | | SAM-dependent methyltransferase responsible for the O-methylation of the hydroxyl group at position 2 of the fucosyl residue of PGL (and possibly p-HBAD) | - | - | |
| Rv2956 | | SAM-dependent methyltransferase responsible for the O-methylation of the hydroxyl group at position 2 of the fucosyl residue of PGL (and possibly p-HBAD) | - | - | |
| Rv2957 | | Fucosyltransferase responsible for the transfer of the third glycosyl residue of the triglycosyl appendage of PGL and p-HBAD | - | - | |
| Rv2958c | | Rhamnosyltransferase responsible for the transfer of the second rhamnosyl residue of the triglycosyl appendage of PGL and p-HBAD | - | - | |
| Rv2959c | | SAM-dependent methyltransferase responsible for the O-methylation of the hydroxyl group at position 2 of the | - | - | |

| | | | | |
|---|---|---|---|---|
| | | rhamnosyl residue linked to the phenolic group of PGL and p-HBAD | | |
| Rv2962c | | Rhamnosyltransferase responsible for the transfer of a rhamnosyl residue onto p-hydroxybenzoic ethyl ester and/or phenolphthiocerol dimycocerosates | - | - |

**Notes**:

- FSD4b-SM contains the *ppsA-E* locus, which is responsible for the synthesis of the phthiodiolone core of the PDIMs, but there are several differences from its *M. tuberculosis* and *M. marinum* counterparts. One important change occurs in the ketoreductase (KR) domains of both PpsB and PpsA PKSs that would cause the corresponding C9 and C11 phthiodiolone glycols to be in the D and L configuration respectively, in contrast to the L and D configuration seen in *M. tuberculosis* and the L and L configuration in *M. marinum* (see Figure S5). According to polyketide synthase biosynthetic logic PpsA of FSD4b-SM should alter the 11-diol stereochemistry to be the same as *M. marinum*, even though FSD4b-SM PpsA has greater amino acid identity to *M. tuberculosis* PpsA.
- FSD4b-SM Mas (mycocerosic acid synthase) has greater similarity to *M. marinum* Mas than to *M. tuberculosis* Mas, including the presence of a conserved tryptophan within the enoylreductase (ER) domain (17). This mutation leads to the production of phthioceranates (also known as mycoceranates) that are added to the phthiodiolone core in *M. marinum* (18). These dextrorotary forms of the mycocerosic acids have all their methyl substituents in the 2 *S* configuration and have only previously been seen in *M. marinum* and *M. ulcerans* (19) (See Figure S5).
- In *M. marinum* and *M. tuberculosis*, the concerted action of a phthiodiolone ketoreductase (Rv2951c) and PDIM methyltransferase (Rv2952c) converts the phthiodiolone dimycocersates (PDIM B) into phthiodiocerol dimycocerosates (PDIM A). However, FSD4b-SM is missing orthologues for both of these genes, suggesting that reduced and methylated PDIMs cannot be formed.
- The substitution of phenolphthiodiolones for phthiodiolones, synthesized by Pks15/1, and their subsequent glycosylation leads to the production of the PGLs (20). Interestingly, *pks15/1* is a pseudogene in *M. tuberculosis* H37Rv, but is intact in *M. tuberculosis* strains of the Beijing lineage, where PGLs are thought to contribute to the hypervirulent phenotype of these strains (21). While *pks15/1* is also intact in *M. canetti* and *M. marinum*, FSD4b-SM is missing this locus, meaning that it cannot produce PGLs.

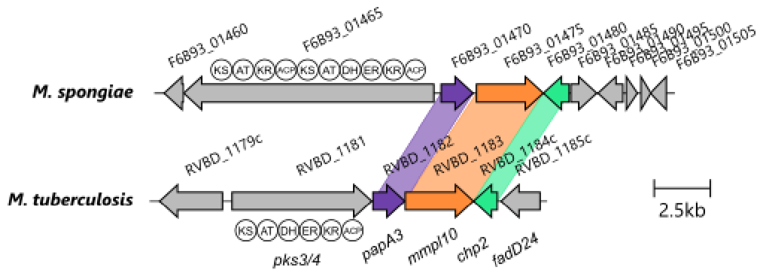

**Figure S4**. Alignment of putative *pks3/4* locus in *M. spongiae* with that from *M. tuberculosis*. PKS domains are indicated underneath each putative PKS protein. Note the absence of a *fadD24* orthologue in *M. spongiae*.

**Table S11.** Orthologues of proteins involved in biosynthesis of trehalose and related compounds.

**Trehalose biosynthesis**

| *M. spongiae* locus tag | Protein | Function | Closest match (locus tag)[a] | % identity | Identified by proteomics |
|---|---|---|---|---|---|
| F6B93_20090 | OtsA/Tps | trehalose-6-phosphate synthase, trehalose biosynthesis | Rv3490 | 87 | Y |
| F6B93_04990 | OtsB2/Tpp | trehalose-6-phosphate phosphatase, trehalose biosynthesis | Rv3372* | 75 | Y |
| F6B93_09900 | TreY | maltooligosyl-trehalose synthase, trehalose biosynthesis | Rv1563c | 75 | Y |
| F6B93_09895 | TreZ | maltooligosyl-trehalose trehalohydrolase, trehalose biosynthesis | Rv1562c | 80 | |
| F6B93_01225 | TreS | trehalose synthase, maltose biosynthesis | Rv0126 | 93 | Y |
| F6B93_21945 | FbpA (Ag85A) | Mycolyltransferase, TDM/mAGP assembly | Rv3804c* | 90 | Y |
| F6B93_11510 | FbpB (Ag85B) | Mycolyltransferase, TDM/mAGP assembly | Rv1886c | 89 | Y |
| F6B93_01240 | FbpC (Ag85C) | Mycolyltransferase, TDM/mAGP assembly | MMAR_0328 | 88 | Y |
| F6B93_01680 | MmpL3 | membrane-associated transporter, flippase, TMM translocation | MKAN_16500 | 82 | Y |

### Sulfolipid and DAT/PAT biosynthesis

| H37Rv locus tag | Protein | Function | *M. spongiae* locus tag | % identity | Identified by proteomics |
|---|---|---|---|---|---|
| Rv0295c | Sft0 | Sulfotransferase, SL-1 biosynthesis | F6B93_02165 | 89 | |
| Rv3822 | Chp1 | membrane-associated acyltransferase, SL-1 biosynthesis/export | F6B93_05155 | 39 | Y |
| Rv3821 | Sap | sulfolipid-1-addressing protein, SL-1 biosynthesis/export | F6B93_14890 | 34 | |
| Rv3823c | MmpL8 | membrane-associated transporter, SL-1 export | F6B93_00620 | 57 | |
| Rv3824c | PapA1 | Acyltransferase, SL-1 biosynthesis | F6B93_01470 | 54 | |
| Rv3820c | PapA2 | Acyltransferase, SL-1 biosynthesis | F6B93_09730 | 55 | |
| Rv3825c | Pks2 | Type I PKS,Elongation of the methyl-branched phthioceranic and hydroxyphthioceranic acids in sulfolipids | F6B93_09705 | 69 | Y |
| Rv3826 | FadD23 | Putative fatty acid AMP ligase, activating fatty acid start units for Pks2 | F6B93_07115 | 66 | |
| Rv0757c | PhoP | sensor kinase, regulation of SL, DAT and PAT biosynthesis | F6B93_19965 | 95 | Y |
| Rv0758c | PhoR | response regulator, regulation of SL, DAT and PAT biosynthesis | F6B93_19960 | 77 | Y |
| Rv1180-Rv1181 | Pks3-Pks4 | Type I PKS, elongation of the methyl-branched mycosanoic and mycolipenic acids found in DAT, TAT, and PAT | F6B93_09725 | 66 | Y |
| Rv1182 | PapA3 | Acyltransferase, PAT biosynthesis | F6B93_01470<br>F6B93_09730<br>F6B93_09715 | 61<br>49<br>49 | |
| Rv1183d | MmpL10 | membrane-associated transporter, PAT export | F6B93_00620 | 64 | |

| | | | | | |
|---|---|---|---|---|---|
| Rv1184c | Chp2 | Acyltransferase, PAT biosynthesis | F6B93_00625 | 50 | |
| Rv1185c | FadD21 | fatty-acyl AMP ligase, PAT biosynthesis | F6B93_07115 | 61 | |
| Rv1660 | Pks10 | Type III PKS, Involved in methyl-brached polyketide biosynthesis in *M. marinum* | F6B93_10340 | 84 | |
| Rv1661 | Pks7 | Type I PKS, Elongation of methyl-branched mycolipenic acids in DAT and PAT | F6B93_10345 | 70 | |
| Rv1662 | Pks8 | Type I PKS, Elongation of methyl-branched mycolipenic acids in DAT and PAT | F6B93_10350 | 65 | |
| Rv1663 | Pks17 | Type I PKS, Elongation of methyl-branched mycolipenic acids in DAT and PAT | F6B93_10350 | 62 | |
| Rv1664 | Pks9 | Type I PKS, Involved in biosynthesis of methyl branched fatty acids | F6B93_10355 | 70 | |
| Rv1665 | Pks11 | Type III PKS, Involved in methyl-branched polyketide biosynthesis in *M. marinum* | F6B93_10360 | 76 | Y |
| Rv2048c | Pks12 | Type I PKS, Involved in the elongation of the alkyl backbone of mycoketides | F6B93_12935 | 79 | |
| Rv3416 | WhiB3 | Regulator of SL, DAT, and PAT synthesis | F6B93_04810 | 93 | |

**LOS biosynthesis**

| ***M. spongiae* locus tag** | **Closest orthologue** | **Protein** | **Function** | **% identity** | **Identified by proteomics** |
|---|---|---|---|---|---|
| F6B93_12950 | MMAR_ 2313 | LosA | glycosyltransferase | 27 | Y |
| F6B93_09730 | MMAR_2343 | PapA4 | polyketide-synthase-associated protein | 76 | |
| F6B93_02315 | MMAR_2309 | Udg | UDP-glucose dehydrogenase | 27 | |
| F6B93_09670 | MMAR_2333 | WcaA | glycosyltransferase | 28 | |
| F6B93_09680 | MMAR_2342 | MmpL12 | putative membrane transporter | 56 | |
| F6B93_09705 | MMAR_2340 | Pks5 | Polyketide synthase | 71 | Y |
| F6B93_09710 | MMAR_2340 | Pks5 | Polyketide synthase | 73 | Y |
| F6B93_09715 | MMAR_2343 | PapA2 | acyltransferase | 51 | |
| F6B93_09720 | MMAR_2343 | PapA2 | acyltransferase | 51 | |
| F6B93_09725 | MMAR_2340 | Pks5 | Polyketide synthase | 72 | Y |
| F6B93_09730 | MMAR_2343 | PapA2 | acyltransferase | 77 | |
| F6B93_09735 | MMAR_2344 | Pks5.1 | Polyketide synthase | 72 | |
| F6B93_09740 | MMAR_2341 | FadD25 | Fatty acyl-AMP ligase | 69 | Y |
| F6B93_09745 | MMAR_2345 | | Carboxymucolonate decarboxylase | 85 | Y |

**Notes**:

- Diacyl trehalose (DAT) and the sulfolipids are acylated forms of trehalose that are restricted to members of the Mtb complex (22, 23).
- Six separate PKS genes (*pks10, pks7, pks8, pks17, pks9* and *pks11*) that encode the methyl-branched fatty acid synthases that are believed to add methyl-branched fatty acids to trehalose as one part of DAT or polyacylated trehalose (PAT) biosynthesis are well conserved in FSD4b-SM (24).
- The proteins responsible for acylating the trehalose moiety, resulting in the formation of DAT, TAT and PAT are encoded by the *pks3/4* locus in *M. tuberculosis* strains, which consists of a polyketide synthase (that is inactivated in the H37Rv

strain, but intact in Erdman and other *M. tuberculosis* isolates) and other related fatty-acid activating and acylating enzymes. While the *pks3/4* locus is well conserved in FSD4b-SM, the *pks3/4* orthologue encodes a protein (FSD4b_00312) that is one whole PKS module longer than Mtb Pks3/4, suggesting the resultant lipid will be 2 carbons longer than the mycolipenolic, mycosanoic or mycolipenic chain usually added to trehalose in the formation of DAT. This structural anomaly would make F6B93_01465 a putative bi-modular iterative PKS, similar to Pks12, which is involved in mycoketide synthesis. Such an enzyme has not previously been reported to provide acyl units for the formation of DAT or more highly decorated trehalose analogues, like TAT and PAT.

- The *pks3/4* locus in FSD4b-SM appears to be missing a critical FadD24 orthologue, which was found to be essential for both DAT and PAT formation in *M. tuberculosis* (25) (Figure S4). This lack of a FadD24 orthologue may explain the lack of detected DAT in our lipidomic analysis. However, given the two modules seen in FSD4b_00312 compared to *pks3/4* it is also possible that a different cellular FadD activates an as yet unrecognised fatty acid for extension and further suggests that the molecule would be a previously unseen acyl-trehalose intermediate.
- Strains that make lipooligosaccharides (LOS) contain a conserved genetic locus analogous to the DAT locus that contains two *pks* genes *(pks5* and *pks5.1*), FadD and Pap orthologues, as well as multiple glycosyltransferases (26–28).
- FSD4b contains a LOS locus, analogous to that in both *M. marinum* and *M. canetti*, however FSD4b *los* also contains unique features that are not seen in other mycobacteria. These include four *pks5* paralogues, compared to two in other species, suggesting a longer acyl chain may be added to the trehalose core. Interestingly, three of the *pks5* paralogues in FSD4b (F6B93_09710, F6B93_09725 and F6B93_09735) are situated next to a corresponding *pap* gene, which encode putative acyltransferases, most likely for transferring acyl chains constructed by each PKS to trehalose, as is seen in sulfolipid, DAT and LOS synthesis in other mycobacteria (29) (Figure S7).
- Compared to *M. marinum*, FSD4b is missing a region encompassing *udgL* to MMAR_2339, which contains several genes responsible for the addition of up to three extra glycosyl units to the LOS core.

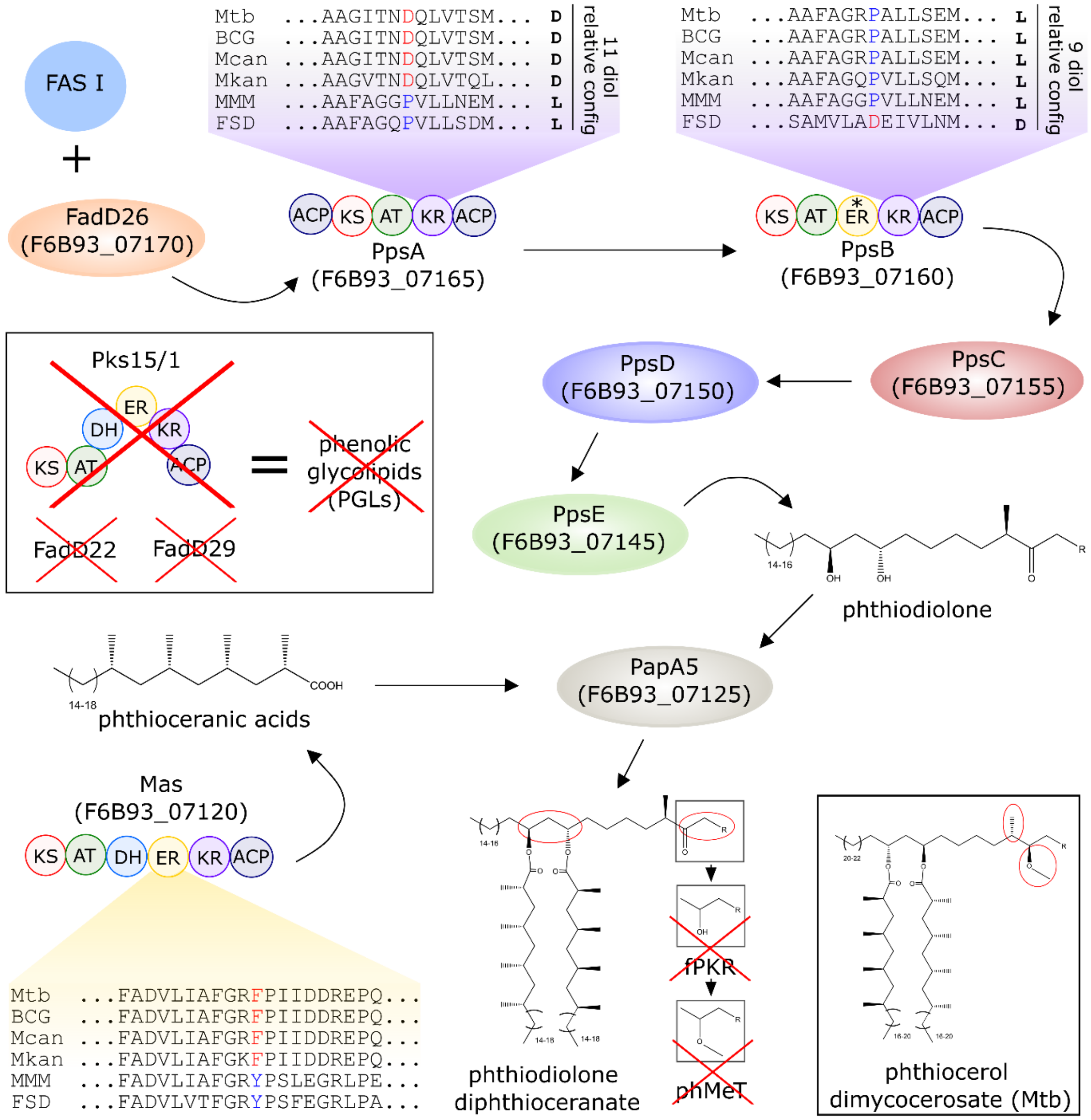

**Figure S5**. Predicted phthiodiolone diphthiocerante biosynthetic pathway in FSD4b-SM and comparison to phthiocerol dimycocerosate biosynthetic pathway in *M. tuberculosis*.

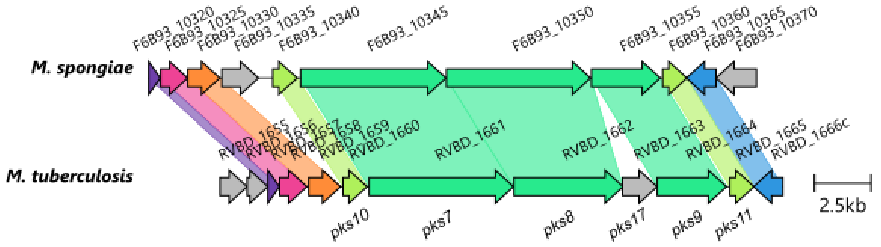

**Figure S6**. Alignment of *pks7-11* region in *M. tuberculosis* with that from *M. spongiae*.

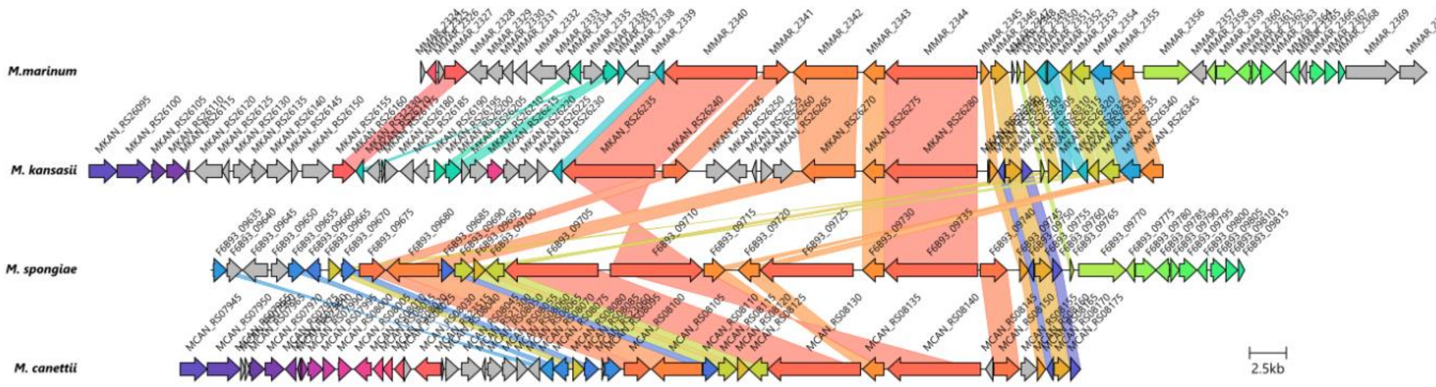

**Figure S7**. Alignment of putative LOS locus in *M. spongiae* with those from other mycobacteria. Note the four *pks* genes in the *M. spongiae* genome.

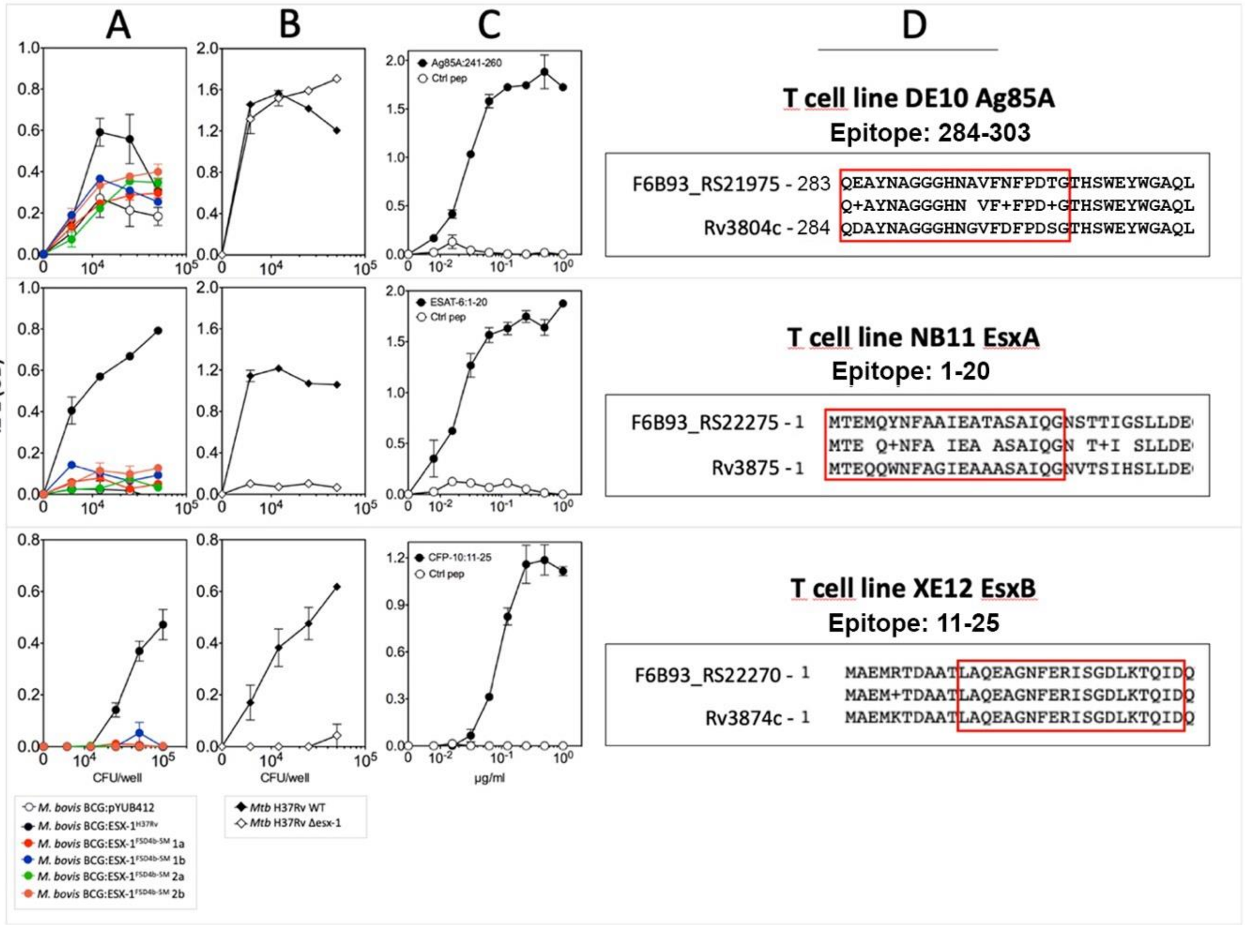

**Figure S8.** Assessment of recombinant *M. bovis* BCG FSD4b-SM Esx-1 protein effector T cell stimulation. (A–D) Measurement of IL-2 production by T cell hybridomas specific for epitopes from *M. tuberculosis* H37Rv antigens Ag85A, EsxA, and EsxB, after co-culture with DCs: (A) infected with various *M. bovis* BCG recombinant ESX-1 strains, (B) infected with WT *M. tuberculosis* H37Rv or an

ESX-1 deletion mutant, or (C) loaded with Ag85A, EsxA, and EsxB peptides encompassing the immunodominant epitopes or control peptide. Shown are concentrations of IL-2 in the co-culture supernatants at 24hr after T cell addition. Error bars are mean and SD of triplicate experiments. (D) Amino acid sequence alignments of *M. spongiae* FSD4b-SM Ag85A, EsxA and EsxB with *M. tuberculosis* H37Rv orthologs, showing conservation and differences with the T cell epitopes.

**Table S12.** Orthologues of proteins involved in the biosynthesis of phospholipids, isoprenoids and related compounds.

| H37Rv locus tag | Protein | Function | *M. spongiae* locus tag | % ID | Identified by proteomics |
|---|---|---|---|---|---|
| Rv0221 | | Putative acyl-CoA:diacylglycerol acyltransferase | F6B93_01770 | 86 | Y |
| Rv0308 | | Putative phosphatidic acid phosphatase | F6B93_02185 | 63 | |
| Rv0436c | PssA | Putative phosphatidylserine synthase | F6B93_03000 | 55 | |
| Rv0437c | Psd | Putative phosphatidylserine decarboxylase | F6B93_03005 | 74 | |
| Rv0534c | MenA | Demethylmenaquinone synthase | F6B93_08210 | 50 | |
| Rv0542c | MenE | o-Succinylbenzoyl-CoA synthase | F6B93_03615 | 79 | Y |
| Rv0548c | MenB | 1,4-Dihydroxy2-naphthoic acid synthase | F6B93_03650 | 93 | Y |
| Rv0562 | | ω,E,E,E-Geranylgeranyldiphosphate synthase | F6B93_03705 | 81 | Y |
| Rv0564c | GpdA1 | Putative glycerol-3-phosphate synthase | F6B93_03725 | 91 | Y |
| Rv0654 | | Carotenoid oxygenase | F6B93_04125 | 78 | Y |
| Rv0895 | | Putative acyl-CoA:diacylglycerol acyltransferase | F6B93_21520 | 42 | Y |
| Rv0989c | | Geranyldiphosphate synthase | F6B93_03705 | 55 | Y |
| Rv1011 | IspE | 4-Diphosphocytidyl-2C-methyl-D-erythritol kinase | F6B93_17860 | 95 | Y |
| Rv1086 | | ω,E,Z-Farnesyldiphosphate synthase | F6B93_17535 | 93 | Y |
| Rv1159 | PimE | Polyprenol phosphomannose-dependent α-1,2-mannosyltransferase | F6B93_17145 | 76 | |
| Rv1411c | LprG | Lipoprotein; putative PIM, LM, and LAM transporter | F6B93_09145 | 80 | Y |
| Rv1425 | | Putative acyl-CoA:diacylglycerol acyltransferase | F6B93_09195 | 89 | Y |
| Rv1551 | PlsB1 | Putative glycerol-3-phosphate acyltransferase | F6B93_09850 | 87 | |
| Rv1760 | | Putative acyl-CoA:diacylglycerol acyltransferase | F6B93_09195 | 35 | Y |
| Rv1822 | PgsA2 | Putative cardiolipin synthase | F6B93_11150 | 78 | |
| Rv2188c | PimB’ | GDP-Man-dependent α-1,6-phosphatidylinositol Mannosyltransferase | F6B93_13570 | 84 | |
| Rv2267c | Stf3 | Putative sulfotransferase | F6B93_20260 | 26 | Y |
| Rv2285 | | Putative acyl-CoA:diacylglycerol acyltransferase | F6B93_14235 | 86 | Y |
| Rv2361c | | ω,E,poly-Z-Decaprenyldiphosphate synthase | F6B93_14935 | 87 | Y |
| Rv2482c | PlsB2 | Putative glycerol-3-phosphate acyltransferase | F6B93_15680 | 89 | Y |
| Rv2483c | PlsC | Putative lysophosphatidate acyltransferase | F6B93_15685 | 89 | Y |
| Rv2484c | | Putative acyl-CoA:diacylglycerol acyltransferase | F6B93_15690 | 90 | Y |
| Rv2524c | Fas | Fatty acid synthetase type I | F6B93_15935 | 91 | Y |
| Rv2610c | PimA | GDP-Man-dependent α-1,2-phosphatidylinositol Mannosyltransferase | F6B93_08585 | 88 | |
| Rv2611c | | Acyltransferase involved in the 6-O-acylation of the Manp residue linked to the 2-position of myo-inositol in PIM1 and PIM2 | F6B93_08580 | 81 | Y |
| Rv2612c | PgsA1 | Phosphatidyl-myo-inositol synthase and/or phosphatidyl-myo-inositol phosphate synthase | F6B93_08575 | 86 | |
| Rv2682c | Dxs | 1-Deoxy-D-xylulose-5-phosphate synthase | F6B93_08280 | 89 | Y |
| Rv2746c | PgsA3 | Phosphatidylglycerophosphate synthase | F6B93_07960 | 80 | |

| | | | | | |
|---|---|---|---|---|---|
| Rv2868c | IspG | 1-Hydroxy-2-methyl-2(E)-butenyl 4-diphosphate synthase | F6B93_07440 | 93 | Y |
| Rv2870c | IspC | 1-Deoxy-D-xylulose 5-phosphate reductoisomerase | F6B93_07430 | 85 | Y |
| Rv2881c | CdsA | Putative CDP-diacylglycerol synthase | F6B93_07390 | 80 | |
| Rv2982c | GpdA2 | Putative glycerol-3-phosphate synthase | F6B93_07005 | 88 | Y |
| Rv3087 | | Putative acyl-CoA:diacylglycerol acyltransferase | F6B93_01770 | 33 | Y |
| Rv3088 | Tgs4 | Putative acyl-CoA:diacylglycerol acyltransferase | F6B93_21535 | 35 | |
| Rv3130c | Tgs1 | Acyl-CoA:diacylglycerol acyltransferase | F6B93_06230 | 73 | |
| Rv3233c | | Putative acyl-CoA:diacylglycerol acyltransferase | F6B93_05615 | 90 | Y |
| Rv3234c | Tgs3 | Putative acyl-CoA:diacylglycerol acyltransferase | F6B93_05615 | 86 | Y |
| Rv3371 | | Putative acyl-CoA:diacylglycerol acyltransferase | F6B93_04995 | 70 | |
| Rv3377c | | Tuberculosinyldiphosphate synthase | F6B93_18415 | 32 | |
| Rv3378c | | Isotuberculosinol synthase | F6B93_01815 | 31 | Y |
| Rv3383c | | ω,E,E,E-Geranylgeranyldiphosphate synthase | F6B93_22300 | 75 | |
| Rv3398c | | ω,E,E-Farnesyldiphosphate synthase | F6B93_22300 | 41 | |
| Rv3480c | | Putative acyl-CoA:diacylglycerol acyltransferase | F6B93_00145 | 82 | Y |
| Rv3581c | IspF | 2C-Methyl-D-erythritol 2,4-cyclodiphosphate | F6B93_20585 | 91 | |
| Rv3582c | IspD | 4-Diphosphocytidyl-2C-methyl-D-erythritol | F6B93_21085 | 73 | |
| Rv3734c | Tgs2 | Putative acyl-CoA:diacylglycerol acyltransferase | F6B93_21520 | 86 | Y |
| Rv3740c | | Putative acyl-CoA:diacylglycerol acyltransferase | F6B93_21535 | 78 | |
| Rv3804c | FbpA | Acyl-CoA:diacylglycerol acyltransferase | F6B93_21945 | 90 | Y |
